# Structural Basis for Nucleobase Activation by the Adenine DNA Glycosylase MutY

**DOI:** 10.64898/2026.01.22.701053

**Authors:** L. Peyton Russelburg, Merve Demir, Karina Cedeno, Sheila S. David, Martin P. Horvath

## Abstract

MutY excises adenine (A) from 8-oxo-guanine:adenine (OG:A) lesions in DNA to initiate base excision repair (BER) and thereby prevent mutations. A catalytic Glu, found at position 43 in the enzyme from *Geobacillus stearothermophilus* (*Gs* MutY), protonates the nucleobase at N^7^ to labilize the N-glycosidic bond. The resulting oxocarbenium ion transition state is stabilized by a covalent DNA-enzyme intermediate and resolved by nucleophilic attack to yield the *beta*-anomer abasic AP site product. The retaining SN1 mechanism for MutY posits deprotonation of the nucleophile by the catalytic Glu. Here we tested kinetic and structural consequences of Glu replacement and found that E43Q and E43S substitution variants were severely impaired, retained measurable activity, but engage the substrate nucleobase in an *anti* conformation, rotated by 180 ° from the *syn* conformation seen in previous substrate complexes. The enzyme-generated AP product is observed in its *alpha*-anomer configuration for these Glu-replacement variants. Comparison with inverting adenine glycosylases that act on RNA or nucleosides shows that MutY’s mechanism is uniquely reliant on one catalytic residue for both leaving group and nucleophile activation, a situation that may serve to ensure only rare adenines paired with OG are excised.

## Introduction

Oxidative damage erodes information stored in DNA. With its low redox potential, guanine reacts readily to generate the most common oxidation product 8-oxo-7,8-dihydroguanine (OG). OG pairs with cytosine but also mispairs with adenine to generate the pro-mutagenic OG:A lesion. The GO repair pathway prevents mutations associated with OG lesions in DNA. An OG-specific base excision repair (BER) DNA glycosylase finds and initiates repair at OG:C lesions by removing the damaged base. In bacteria this OG DNA glycosylase is Fpg encoded by the MutM gene,^[1,2]^ while eukaryotes and archaea encode a functional ortholog called OGG1.^[3–5]^ MutT (MTH1 in humans) hydrolyzes OG nucleotide triphosphates, thus minimizing OG incorporation into the DNA daughter strand.^[6,7]^ MutY and its homolog MUTYH in mammals provide the last line of protection. These adenine DNA glycosylases intercept OG:A lesions and initiate BER by cleaving the N-glycosidic bond of the mismatched 2’-deoxyadenosine.^[8]^ Although MutY will process G:A mispairs *in vitro* and is often annotated as an A/G specific adenine DNA glycosylase, its primary substrate in cells is the OG:A lesion as evidenced by genetic complementation and cellular repair assays.^[9–11]^

The importance of the GO repair pathway is highlighted by consequences when the system underperforms in cancer patients. For example, biallelic defects in the human gene encoding MUTYH lead to a high rate of G:C → T:A mutation in somatic cells, eventually disabling tumor suppressor genes to explain early onset cancer, first recognized for a pattern of inherited predisposition for colon polyps and termed MUTYH associated polyposis (MAPS).^[8,12,13]^

MutY/MUTYH belong to the Helix-hairpin-Helix (HhH) DNA glycosylase superfamily which includes EndoIII and OGG1. As is true for most DNA glycosylases, EndoIII and OGG1 find and initiate repair by removing a damaged base. Unique among these glycosylases, MutY excises a chemically normal purine and must integrate information from the partner strand to distinguish appropriate and rare substrate adenines paired with damaged OG from other, much more abundant adenines in DNA. MutY employs OG-specific interactions provided by three residues that comprise the FSH motif in its C-terminal domain (CTD). The FSH motif establishes two hydrogen bonds with OG that are not made with G,^[14]^ is highly conserved across enzymes from bacteria and eukaryotes,^[15,16]^ and replacement of these FSH residues disables mutation suppression function in bacteria.^[14,17]^

All HhH DNA glycosylases employ a catalytic Asp residue to facilitate N-glycosidic bond cleavage. MutY, but not other HhH family members, complements the catalytic Asp with a catalytic Glu which features prominently in the first step of a dissociative SN1 mechanism.^[18,19]^ Replacement of the Asp with Asn, and replacement of the Glu with Gln or Ser inactivated MutY from *Escherichia coli* (*Ec* MutY), explaining increased mutational burden in cultures and an inability to repair synthetic OG:A lesions in cells.^[20,21]^ Notably, replacement of the Asp with Glu, or Glu with Asp, retained some activity, confirming the importance of a carboxylic acid at both positions. Non-synonymous mutations in human MUTYH that replace these critical residues are associated with pathology in cancer patients as documented with ClinVar^1^.^[22]^

Further evidence for the importance of the catalytic Asp and Glu comes from X-ray crystal structures. The fluorinated lesion recognition complex (FLRC) structure showed Glu43 in MutY from *Geobacillus stearothermophilus* (*Gs* MutY) within H-bonding distance of N^7^ of the substrate adenine base,^[19]^ thereby identifying the Glu as the acid/base catalyst in early steps of the mechanism. Proton transfer from Glu to adenine pulls electrons from the N-glycosidic bond, thereby activating the nucleobase as a leaving group (**Figure 1**). The N-glycosidic bond breaks and reforms reversibly before irreversible water hydrolysis results in the apurinic/apyrimidinic (AP) site product.^[18]^ The transition state analog complex (TSAC) structure of MutY revealed close approach of both Asp144 and Glu43 with the 1N group ((3R,4R)-4-(hydroxymethyl) pyrrolidin-3-ol) that mimics the shape and charge of the oxocarbenium ion transition state, consistent with electrostatic stabilization.^[23]^

**Figure 1.**
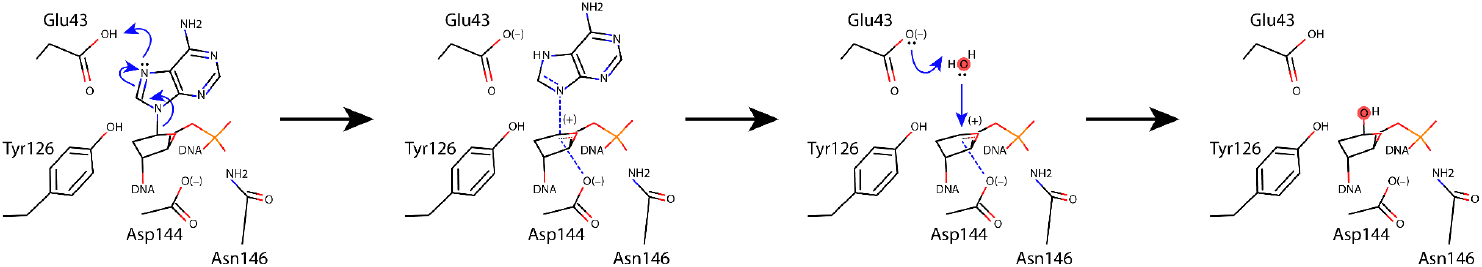
Mechanism of MutY. A glutamic acid (Glu43) initiates glycosidic bond cleavage by protonating N^7^ of adenine and guides stereochemistry of nucleophile attack in the retaining mechanism for MutY. Drawn with Avogadro v1.2.0,^[24]^ on the basis of structures of the FLRC (PDB ID 3g0q),^[19]^ the TSAC (PDB ID 6u7t),^[23]^ and structures of *Gs* MutY (N146S) in complex with the enzyme-generated AP site.^[25]^

The crystal structures indicated different points of attack for the hydrolysis phase of the mechanism. The FLRC inhibitor-enzyme structure showed electron density for a solvent molecule on the 3’ face, close to Asp144 that was interpreted as the nucleophile.^[19]^ However, the TSAC transition state structure showed that this 3’-face water is an unlikely nucleophile because it is far away (4 Å) from the C^1’^ position and Asp144 blocks its path.^[23]^ In the TSAC, a water held in place by Glu43 was much closer (3 Å), with unhindered access to the electrophile from the 5’ face.^[23]^ This 5’-face water was not evident in the FLRC structure because the nucleobase was still present. The product of enzyme-catalyzed methanolysis accumulated with retention of stereochemistry, further supporting attack from the 5’ face.^[23]^ Structures for *Gs* MutY (N146S) complexed with enzyme-generated AP product confirmed the *beta* stereoisomer expected for a double-displacement, retaining mechanism.^[25]^ The double displacement, retaining mechanism for MutY predicts a covalent DNA-enzyme intermediate, which has been observed through QM/MM calculations.^[26]^

The MutY mechanism implicates the catalytic Glu in both early and late steps.^[23,25]^ Herein, we replaced Glu43 in *Gs* MutY with two different, catalytically inert residues, Ser and Gln, evaluated the E43S and E43Q variant enzymes in glycosylase assays that revealed impaired glycosylase activity, and determined structures for substrate and enzyme-generated AP product complexes. The overall architecture of MutY was preserved, however the details at the active site were different and revealing. We captured a substrate complex with the adenine base oriented in an unexpected *anti* conformation; all previous structures show the substrate base in a *syn* conformation. A structure of the enzyme-generated AP product in complex with E43S *Gs* MutY revealed the *alpha*-anomer, suggesting an inverting mechanism contributes to product accumulation when the catalytic Glu is unavailable. These results point to a previously unrecognized role for the catalytic Glu in determining the pose of the nucleobase and are consistent with nucleophile activation by the catalytic Glu in the final step of the retainingmechanism for MutY. The multiple roles played by the catalytic Glu in both leaving group and nucleophile activation make MutY’s mechanism unique among adenine N-glycosylases.

## Results and Discussion

### Glycosylase activity for E43S and E43Q *Gs* MutY

We previously evaluated the kinetics for wildtype *Gs* MutY under single- and multiple-turnover conditions.^[23]^ Kinetic **Scheme 1** applies for this enzyme that is subject to product inhibition.^[27]^ *K*_*d*_ characterizes the rapid equilibration of bound and unbound DNA, *k*_2_ represents the rate of chemical conversion from substrate to product, and *k*_3_ is the rate of product release, which is the limiting step under multiple turnover conditions.^[27]^ Here, we measured and compared reaction rates for Glu43 replacement variants of *Gs* MutY, monitoring the progress by gel electrophoresis to separate radiolabeled substrate and product DNAs. Accumulation of product catalyzed by the E43S and E43Q variants of *Gs* MutY showed one phase, indicating a step prior to product release was now rate limiting for these impaired enzymes (**Figure 2**). Accordingly, we applied a simpler analysis as described previously,^[27]^ to determine that adenine excision rates (*k*_2_) were 0.015 (± 0.004) min^-1^ and 0.0098 (± 0.0004) min^-1^ for the E43Q and E43S enzymes, respectively. Replacement of Glu43 with Gln and Ser slowed the reaction by a factor of 3,600 and 5,500 relative to wildtype *Gs* MutY, previously characterized with *k*_2_ = 54 (± 4) min^-1^.^[23]^

**Scheme 1.**
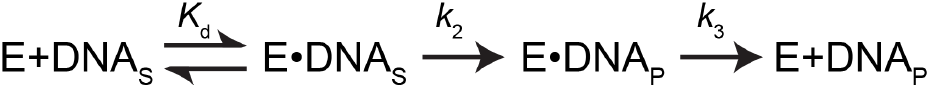
Kinetic scheme for MutY. Kd is rapid equilibration of bound and unbound substrate DNA. For the wildtype enzyme with slow product release, *k*_3_ < *k*_2_, burst kinetics with two phases are observed. If adenine excision, *k*_2_, becomes rate-limiting, one phase is observed.

**Figure 2.**
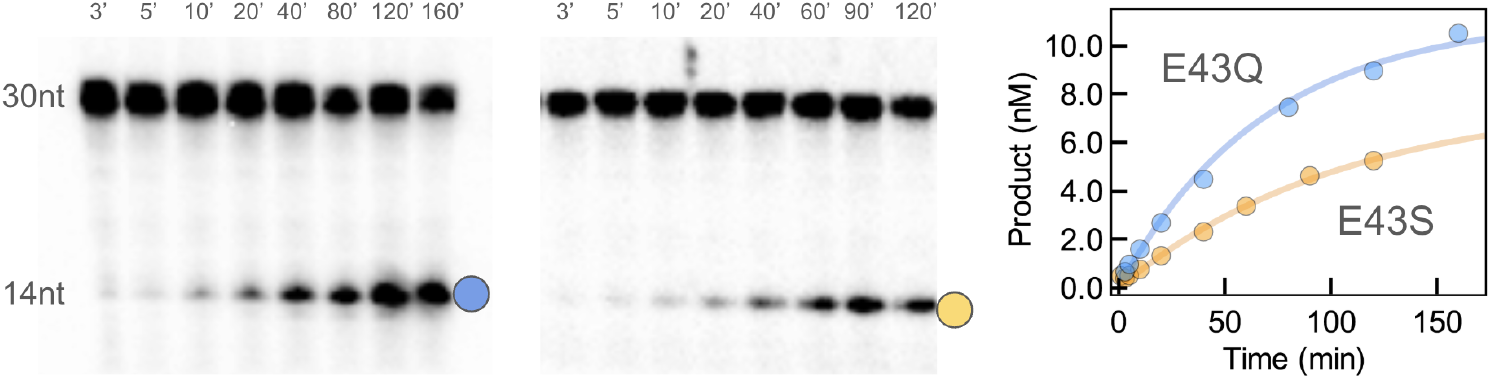
Adenine glycosylase activity for E43Q and E43S *Gs* MutY. DNA containing OG across from adenine was combined with enzyme, incubated at 60 °C, and reactions were quenched by addition of sodium hydroxide to cleave DNA at the AP product. Bands correspond to the low-mobility substrate DNA (30 nt) and high-mobility product DNA (14 nt) and were visualized by autoradiography. Time course data were fit to a single exponential phase. Trials were repeated 3 times (E43S) or 5 times (E43Q).

Replacement of catalytic residues reveals their importance by comparison of the variant and wildtype enzymes.^[28–31]^ Our previous work showed that replacement of this catalytic residue in MutY from *E. coli* resulted in a catalytically inactive enzyme that was unable to repair OG:dA lesions in a cellular repair assay.^[20]^ Although impaired, *Gs* MutY enzyme variants with Ser and Gln substituting for the catalytic Glu retained activity. As *Gs* MutY comes from an extreme thermophile, assays described in the current work were at 60 °C over two hours, conditions that are not feasible for *Ec* MutY, an enzyme from a mesophile. Activity remaining for the E43S and E43Q variants of *Gs* MutY is significantly greater than the activity for variants of subtilisin with catalytic triad residues replaced by Ala, which are 19,000-fold to 1.3 million-fold slower than the wildtype enzyme.^[32]^

### X-ray crystallography

To better understand mechanistic contributions of the catalytic Glu and the source of significant residual adenine glycosylase activity, we crystallized the E43S and E43Q variants in complex with DNA containing OG paired with adenine (dA) or purine (dP) and determined the structures by crystallography. Diffraction data with synchrotron generated X-rays were measured to high resolution and the structures were determined by molecular replacement followed by model rebuilding and refinement. Importantly, the substrate base was omitted from initial refinement so as to observe unbiased discovery maps, which revealed intact substrate for some structures and conversion to the AP (abasic site) product in other cases.

We selected for complete model refinement three representative structures on the basis of data quality (**Table 1**), and will refer to these structures by the replacing residue at position 43 and the nature of the DNA lesion. For example, E43S_OG:dA_ indicates the E43S variant of *Gs* MutY in complex with DNA containing OG across from adenine. Likewise, E43Q_OG:dP_ denotes the E43Q variant with an OG:purine (OG:dP) lesion, and E43S_OG:AP_ is the structure of E43S *Gs* MutY in complex with the enzyme-generated apurinic/apyrimidinic (AP) site. In other contexts AP refers generally to any abasic linker within DNA, often the tetrahydrofuran linker, which is a synthetic component not found naturally in DNA; however, here AP signifies the authentic product of the MutY-catalyzed reaction. After multiple rounds of energy minimization interleaved with model rebuilding, the average temperature factors for these structures ranged from 50 to 58 Å^2^, typical for DNA-MutY complexes, and R-free and R-work values ranged from 22-25% (R-free) and 19-22% (R-work). The final refined models were characterized by high-quality expectations for Ramachandran plots, clash avoidance, and bond geometry (see Supplementary Information **Table S1**).

**Table 1.** Structures for Glu43 replacement variants of *Gs* MutY.

|  | E43S<sub>OG:dA</sub> | E43Q<sub>OG:dP</sub> | E43S<sub>OG:AP</sub> |
| --- | --- | --- | --- |
| Position 43 | Ser | Gln | Ser |
| DNA active site | Adenine | Purine* | AP product |
| Resolution (Å) | 2.60 | 2.20 | 1.68 |
| R-free | 0.253 | 0.234 | 0.245 |
| R-work | 0.208 | 0.192 | 0.222 |
| PDB ID | 8dwf | 8dwe | 8dwd |

\* Partial occupancy

### Overview of structures

Two structures, E43S_OG:dA_ and E43Q_OG:dP_, were obtained in the P2_1_ space group with two copies of the MutY-DNA complex in the asymmetric unit (**Figure 3A** and **Figure 3B**). This space group arises for some of our unpublished structures for *Gs* MutY, and has been reported previously for *Gs* MutY complexed to its anti-substrate,^[33]^ and the N146S variant of *Gs* MutY in complex with DNA containing a transition state analog.^[25]^ A third structure, E43S_OG:AP_, crystallized in the P2_1_2_1_2_1_ space group (**Figure 3C**), which has been more commonly observed for *Gs* MutY structures. The double stranded DNA could be built into robust electron density for ten complete base pairs, including the intra-helical OG nucleotide across from its extra-helical partner. Weaker density clouded the view for some of the 5’-overhanging nucleotides and these remained unpaired even in the P2_1_ space group where 10-mer duplexes stack end-to-end at the non-crystallographic 2-fold axis (**Figure 3A** and **Figure 3B**).

**Figure 3.**
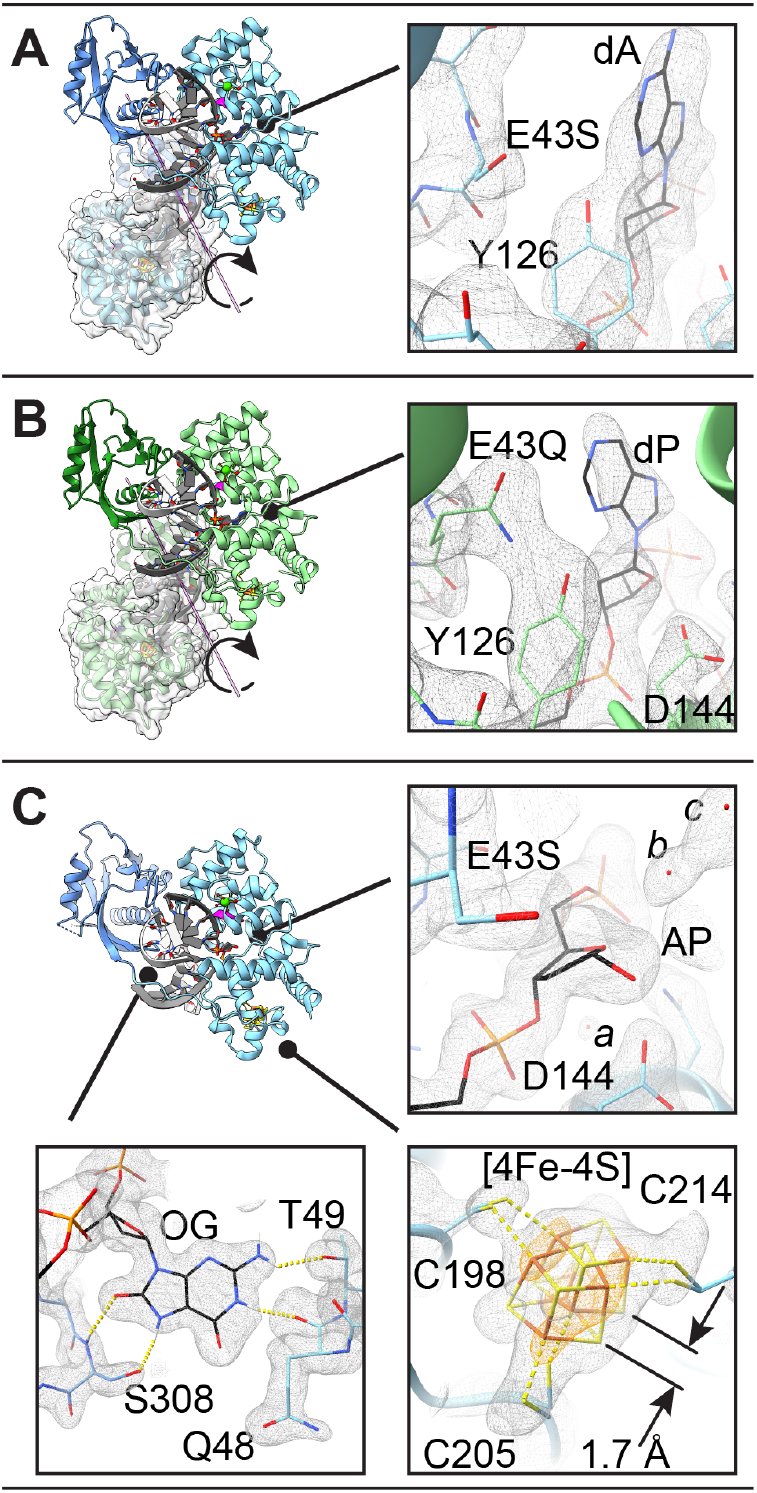
Structures of *Gs* MutY with Glu43 replacement. DNA is colored grey and black. The site of replacement is highlighted with magenta in the ribbon overviews. **A-B.** E43S_OG:dA_ (**A**) and E43Q_OG:dP_ (**B**) crystallized in the p2_1_ space group with two DNA-MutY complexes in the asymmetric unit. Detailed views show the active site with simulated annealing composite omit maps (grey) contoured at the 1-*σ* r.m.s.d. level. **C.** E43S_OG:AP_ crystallized in the p2_1_2_1_2_1_ space group with one DNA-MutY complex in the asymmetric unit. Detailed views show the active site, the iron-sulfur metal center, and the OG recognition site, with the simulated annealing composite omit map contoured at the 1.2-*σ* level and the anomalous map contoured at the 6-*σ* level (orange). Note the alternate conformations for the [4Fe-4S] metal center and *alpha* stereoconfiguration for AP product (**C**). Solvent molecules are found nearby the AP product and Asp144 (labelled *a, b, c*).

The overall structures are comparable to those previously described for MutY complexed to DNA. The enzyme is crescent shaped with two domains connected by a linker nearly encircling the DNA. An iron-sulfur cluster [4Fe-4S] is chelated by four Cys residues in the N-terminal domain (NTD). For the E43S_OG:AP_ structure captured at relatively high resolution (1.68 Å) two alternate conformations are distinguishable in anomalous difference maps for the [4Fe-4S] metal center (**Figure 3C**), a feature noted in other high-resolution views of *Gs* MutY with amino acid replacements.^[25,34]^

Strong electron density defined the replacing residue at position 43 and the nature of the DNA at the active site (**Figure 3**). The NTD inserts intercalating wedge residues Gln48 and Tyr88 into the minor groove to bend the DNA at an angle of ∼50 degrees. The substrate base (**Figure 3A** and **Figure 3B**) or AP product (**Figure 3C**) are extruded into the active site defined by residues Tyr126, Asp144, and Asn146 along with the replacing residue at position 43. Interactions with the orphaned, yet intra-helical OG base in these structures were preserved as described for previous structures.^[14,23,35]^ Figure 3C shows a representative detailed view of the OG lesion as found in the E43S_OG:AP_ structure. The peptide carbonyl group of Gln48 and the side chain of Thr49 make hydrogen bonds with N^1^ and N^2^ along the Watson-Crick-Franklin face of OG. The C-terminal domain (CTD) contributes the FSH motif at the tip of a *beta* loop,^[14]^ with Ser308 making OG-specific hydrogen bonds with N^7^ and O^8^ found along the Hoogsteen face which distinguishes OG from undamaged guanine.

### Enzyme activity by crystallography

The three structures selected for complete refinement were representative of several crystals with the same substrate *versus* product status. In other words, we have reproducible results applying X-ray crystallography to diagnose progress of the enzyme-catalyzed glycosylase reaction. Outcomes as observed in electron density maps obtained from multiple crystals of each class are summarized in **Table S2**. Substrate was consistently observed in four crystals for the E43S variant complexed with DNA containing dA across from the OG lesion. Electron density was missing for the base moiety in seven examples for the E43S variant originally complexed with DNA containing purine across from the OG lesion. The three crystals measured for E43Q complexed with the OG:dP lesion consistently revealed density for the base moiety, albeit with weaker signal in |Fo| - |Fc| difference maps, indicating partial conversion to product. The group occupancy for the purine base in the fully refined E43Q_OG:dP_ structure was 0.7-0.8. Thus, crystallography corroborates our finding that Glu43 replacement impairs but does not completely ablate glycosylase activity.

The presence of substrate base or AP product in crystals likely reflects multiple processes including enzyme catalysis and crystal growth. Divalent metal ions such as Ca^2+^ included with crystallization reactions inhibit *Gs* MutY and *Gs* MutY (N146S).^[25]^ To encourage intermediates and products, DNA and enzyme were combined and incubated at ambient temperature for 30-90 min or at 4 °C for several hours before adding crystallization solutions with calcium acetate (**Table S5)**. We speculate that slow enzyme catalysis favors crystals with the substrate base and also crystals with the AP product. Rapid catalysis may have prevented growth of crystals, perhaps due to AP product instability. This model explains why the E43Q enzyme, which is faster than E43S (**Figure 2**), could not be crystallized with DNA containing dA across from OG, the substrate that is processed faster than dP (**Figure S1**), and why the wildtype enzyme failed to crystallize with either substrate.

For the E43S enzyme and DNA with the OG:dP lesion, catalysis and crystallization apparently collaborated to generate the enzyme-bound AP site product trapped in crystals. DNA with OG across from dP was a poor substrate in glycosylase assays for E43S and E43Q (**Figure S2**). With only 50% conversion to product by E43S acting on OG:dP during overnight incubation at 60 °C, preincubation of enzyme and DNA for 30 min at ambient temperature would not be sufficient to accumulate significant product from the dP substrate. To explain 100% AP product in the crystals of E43S_OG:AP_, metal ions must have failed to inhibit the E43S variant enzyme so that catalysis proceeded concurrently with and possibly after crystallization. In support of this model, enzyme activity has been demonstrated *in crystallo* for *Gs* MutY (N146S).^[25]^

### Active site structure for substrate

To structurally evaluate the contribution of Glu43 we focused attention on the active site of MutY looking for elements that were preserved and things that changed in response to replacement of Glu43. For the Ser substituted variant with adenine in the active site, E43S_OG:A_, we observed tight fitting electron density for a small residue at position 43 (**Figure 3A**), consistent with Ser replacing Glu. Previous structures of wildtype *Gs* MutY, complexed with DNA containing a non-cleavable substrate analog,^[19]^ showed Glu43 interacting with Tyr126 and engaged with N^7^ of the 2’-fluoro-2’-deoxy-adenine (**Figure 4A**). Relative to this reference structure, Ser43 shows a substantially different and reduced set of interactions (**Figure 3A** and **Figure 4B**). Tyr126 has shifted to partly fill the space vacated by Glu43 replacement. Other active site residues preserved interactions seen in previously described structures. For example, catalytic residues Asp144 and Asn146 maintained hydrogen bonding interactions with each other and the DNA backbone. From this we may conclude that perturbations consequent to E43S replacement were fairly localized.

**Figure 4.**
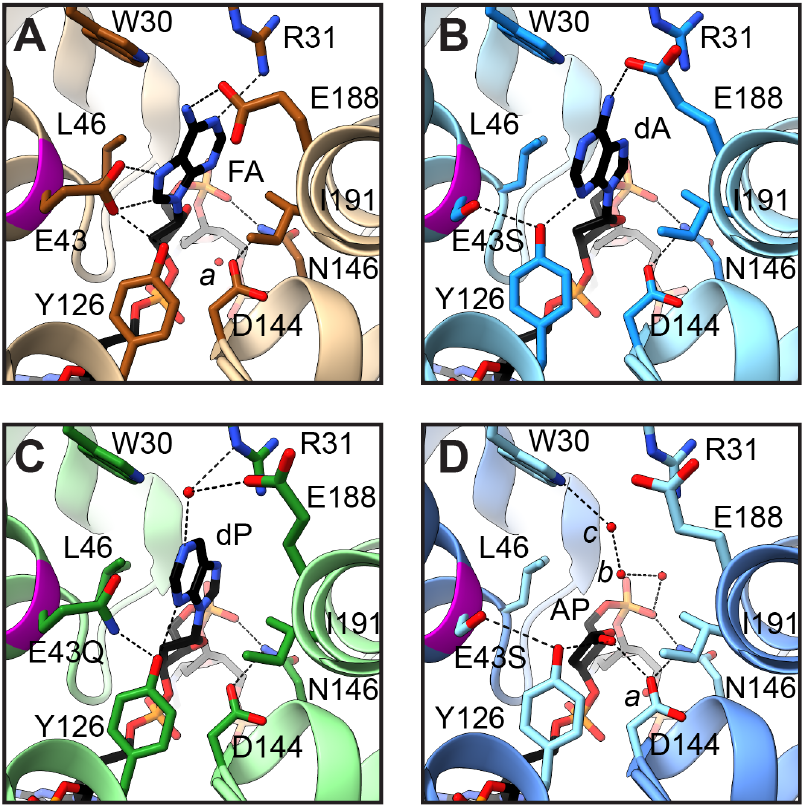
Active site structures. **A.** The nucleobase of 2’-fluoro-2’-deoxy-adenosine (FA) orients in a *syn*-like conformation (Chi-1 = -30.5°) and hydrogen bonds with Glu43 and Glu188 in the active site of the FLRC (PDB ID 3g0q). **B.** Adenine (dA) orients in an *anti*-like conformation (Chi-1 = -172°) and hydrogen bonds with Tyr126 and Glu188 in the active site of the E43S variant enzyme. **C.** Purine (dP) orients in the *anti*-like conformation (Chi-1 = -186°) and hydrogen bonds with Tyr126 in the active site of the E43Q variant enzyme. For each of these substrate complexes the nucleobase is sandwiched between Leu46 and Ile191. **D.** The AP product, generated by E43S acting on DNA with dP, is observed as the ring-closed *alpha* anomer. Solvent molecules nearby the AP site are labelled (*a, b*).

Local perturbations to DNA structure are evident. The adenine base adopted an *anti*-like orientation (**Figure 3A** and **Figure 4B**), with O^4’^ and C^4^ in *trans* and rotated by ∼180° relative to the *syn*-like conformation observed for prior substrate complexes, including the lesion recognition complex (LRC),^[35]^ the fluoro-A (FA) lesion recognition complex (FLRC),^[19]^ and a recently described structure for the N146S variant of *Gs* MutY complexed with purine (N146S_OG:dP_).^[25]^ Evidence for the unexpected *anti* conformation comes from maps calculated after initial molecular replacement, which outlined a nucleobase with an exocyclic group (Supplementary Information **Figure S2**). Replacement of Glu43 and rotation of the N-glycosidic bond change the hydrogen-bonding interactions available to the nucleobase. Glu43 of the wildtype enzyme approaches closely with N^7^ and C^8^ of FA in the *syn*-like conformation (**Figure 4A**).^[19]^ In the *anti* conformation, N^7^ fails to find a partner and N^3^ of dA hydrogen-bonds with the hydroxyl group of Tyr126. Glu188 adapts its rotamer configuration to preserve the hydrogen bond with exocyclic N^6^ (**Figure 4B**). Switching from *syn* to *anti* is coupled with a different sugar pucker: C^2’^-*endo* for *syn* becomes C^3’^-*exo* for *anti*. The net effect preserves the position of the nucleobase, sandwiched between Leu46 and Ile191, for the wildtype enzyme and for E43S_OG:dA_ described here.

We also observed the *anti*-like conformation with C^3’^-*exo* sugar pucker for the dP nucleobase in the E43Q_OG:dP_ structure (**Figure 3B** and **Figure 4C**), an outcome that indicates the *anti* conformation does not rely on N^6^ interactions that are available for dA but absent for dP. We also infer that, in the absence of Glu43, the *syn*-like orientation is unstable relative to the *anti* orientation. In the E43Q_OG:dP_ structure, Gln43 hydrogen bonded with Tyr126 and adopted the same rotamer as seen for Glu43, meaning it was in the correct position to interact with a *syn* nucleobase. However, the dP nucleobase remained disengaged from Gln43 by rotation about the N-glycosidic bond. We may conclude that shifting the *anti-syn* equilibrium toward the *syn* conformation is coupled with the strongly shared proton connecting glutamic acid of the wildtype enzyme and the adenine nucleobase.

Glutamic acid at position 43 is understood to be the acid/base catalyst that initiates adenine excision by protonation of N^7^ (**Figure 1**).^[19]^ We examined the structures looking for other residues that could serve as an alternate acid/base catalyst in the E43S and E43Q variant enzymes. R31 and E188 make a solvent-mediated interaction with N^1^ of dP in the structure of E43Q_OG:dP_. Additionally, Tyr126 hydrogen bonds with N^3^ of the nucleobase in the structures for E43S_OG:dA_ and E43Q_OG:dP_. Proton transfer from the water molecule held in place by R31 and E188 to N^1^ or from Tyr126 to N^3^ may thus contribute to impaired glycosylase activity measured for these variant enzymes (see **Figure S3**). In support of this idea, the wildtype enzyme from *E. coli* (*Ec* MutY) excised a hydrophobic isostere of adenine that lacked N^7^ and retained N^3^, 7-deaza-adenine,^[36]^ and the isostere lacking N^3^, 3-deaza-adenine, was excised at a rate 150-fold slower relative to the authentic substrate,^[10]^ observations that highlight the contribution of N^3^ for adenine excision.

### Structure for enzyme-generated product

Electron density for a substrate nucleobase moiety was missing in the E43S_OG:AP_ structure. Absence of electron density for the purine base in the discovery maps calculated for initial rounds of refinement, well defined simulated annealing composite omit maps (**Figure 3D**), and difference maps calculated for the final model refined with data to the 1.68 Å resolution limit (**Figure S4**) provide strong evidence that the base has been excised. The detached free base was evidently not retained following cleavage of the N-glycosidic bond since unexplained volumes of electron density were absent in neighboring regions. Electron density surrounding the abasic site defined an AP group in its closed-ring furanose form, with density extending beyond C^1^’ in an equatorial disposition, consistent with a hydroxyl group in the *alpha* stereoconfiguration (**Figure 3D** and **Figure S4B**) and very different from the *beta* stereoconfiguration observed for the AP product generated by the N146S enzyme (**Figure S4A**).^[25]^

We took special care in selecting the C^4’^-*endo* sugar pucker and *alpha* stereoisomer for the AP site. The *alpha* anomer with C^4’^-*endo* pucker appeared to best fit electron density best (**Figure 3D**), refined with reasonable temperature factors for all atoms in the AP group, and avoided steric clashes (Supplementary Information **Table S7**). The *alpha* anomer with C^4’^-*endo* pucker places the O^1’^ group within hydrogen bonding distance of Asp144 and Tyr126 (**Figure 4D**) and was unexpected since previous studies show *Gs* MutY with Glu at position 43 converts substrate to the *beta* anomer in both methanolysis and hydrolysis reactions.^[23,25]^ Comparison of the E43S_OG:AP_ structure described here with N146S_OG:AP_, described previously,^[25]^ indicates Glu43 replacement has dramatically altered interactions with the AP product (**Figure S4**), and suggests an alternative inverting mechanism may contribute to product accumulation (**Figure S3**).

### Implications for MutY’s catalytic mechanism and regulation

Our work reinforces and extends the idea that recruitment of a catalytic glutamic acid residue was a cornerstone innovation for the evolution of an OG:A-specific adenine glycosylase with a retaining mechanism. DNA glycosylases are tasked with finding and precisely identifying rare substrates among vast amounts of normal DNA, placing evolutionary pressure to finetune substrate specificity. The search for substrate while avoiding off-target decoys is made especially challenging for MutY which excises an undamaged adenine nucleobase. Most other glycosylases remove a damaged base, and therefore can take advantage of distinct chemical groups for recognition and the inherent instability of the N-glycosidic bond following chemical modification.^[37–40]^ In the absence of chemical damage, excision of the adenine base places additional demands for catalysis and substrate recognition. These demands likely contributed to the recruitment of a second catalytic carboxylic acid residue. All HhH glycosylase family members employ an Asp for catalysis (Asp 144 in Gs MutY), and MutY augments this with the catalytic Glu that is found only among MutY, and closely related MAG1 / MIG enzymes.^[16]^

Since the nucleobase being excised is highly prevalent, MutY must strictly integrate information from the OG-recognition site to distinguish appropriate OG:dA substrates. Off target activity would lead to massive AP accumulation and genome instability. This molecular logic makes clear the need for an “*on hold*” state that precedes the fully licensed catalytic state of MutY. Switching between states likely involves allosteric communication among distant sites, including the OG-site, the active site, and the [4Fe-4S] metal center. Connectivity with the metal center is supported by recently reported new structures for *Gs* MutY and human MUTYH complemented by molecular dynamic analysis.^[34]^ In that work, variants of MutY that interrupt the connection, namely N146S and R141Q, retain structure, bind substrate DNA normally, yet are catalytically disabled.^[34]^ The structures for E43 replacement variants of MutY described herein extend these ideas and suggest that an “on hold” state could be created (and released) by taking advantage of *anti-syn* conformational switching. With the nucleobase in an *anti*-like conformation, Glu43 would be disengaged from N^7^, thereby keeping catalysis in check. In this model, switching to a *syn*-like conformation would release the “on hold” state, allow catalytic engagement between glutamic acid and the nucleobase, and initiate efficient adenine excision.

### The covalent intermediate

The retaining mechanism of MutY predicts a covalent DNA-enzyme intermediate,^[23,25]^ and a covalent crosslink from C^1’^ in DNA to the carboxylate of Asp 144 in MutY has been observed in QM/MM structures.^[26]^ The hope of capturing this covalent DNA-MutY intermediate in a crystal structure provided impetus for exploring Glu43 replacement variants. We reasoned that purine, with its enhanced leaving group potential, would increase the rate of forming the covalent intermediate, while Glu replacement would slow its destruction. Similar Glu replacement with Gln led to capture of covalent enzyme-glycosyl intermediates for hen egg white lysozyme and the human nucleotide salvage factor DNPH1.^[41,42]^ However, expectations were not fulfilled for our MutY system. Substrate purine was processed by E43S and E43Q enzymes only very slowly (**Figure S1**), suggesting that purine is a poor leaving group for the enzyme-catalyzed reaction, perhaps consequent to adopting the *anti* configuration. The “on hold” state proposed for regulating MutY catalysis may further explain why the expected covalent intermediate of MutY eluded capture. Release of the “on hold” state presents additional kinetic steps preceding catalysis. If any of these earlier steps are slower than the final nucleophile attack, the MutY covalent intermediate will remain poorly populated.

### Catalytic strategies for adenine glycosylases

A comparison of MutY with other DNA glycosylases and O-glycosidases has been discussed previously.^[23]^ Here we compare and contrast the dsDNA adenine glycosylase MutY with other N-glycoside bond-hydrolyzing enzymes acting on adenine in the context of structured RNA and ribonucleoside substrates, namely the ribosome inactivating proteins (RIPs) ricin A-chain and saporin-L3, and the purine-specific nucleoside hydrolase (NH) from *Trypanosome vivax* (**Figure 5**). These enzymes each solve the challenge of breaking a *beta*-N-glycoside bond connecting C^1’^, of D-ribose or 2’-deoxyribose, to N^9^ of adenine, as well as other purine substrates. As the proteins have completely different folds, the solutions are the result of convergent evolution. Kinetic isotope effect (KIE) analysis demonstrated differences in transition state structure,^[18,43–45]^ yet each adenine glycosylase breaks the N-glycoside bond in a dissociative SN1-like mechanism, leading to a neutral or positively charged leaving group and a positively charged oxocarbenium ion intermediate that is resolved by electrophile migration to an activated nucleophile. The strategies employed for leaving group stabilization are distinct in detail but consistently involve multiple hydrogen bonds to the nucleobase as inferred from the substrate ground state structure in the case of MutY,^[19]^ and from structures of transition state mimics complexed with ricin A-chain, saporin-L3 and NH.^[46,47]^ The nucleophile is activated by a carboxylate group, either Asp or Glu, with attack from the *beta* face distinguishing MutY from the RNA and nucleoside glycosylases, which resolve the transition state with attack from the *alpha* face.^[23,25,46,47]^ These adenine glycosylases treat the oxocarbenium ion transition state intermediate very differently, with MutY providing strong covalent interactions,^[23,25,26]^ and the other glycosylases avoiding covalent catalysis. As a result of these differences, the abasic AP product retains stereoconfiguration in the case of MutY and inverts stereoconfiguration in the case of RIPs and NH.

**Figure 5.**
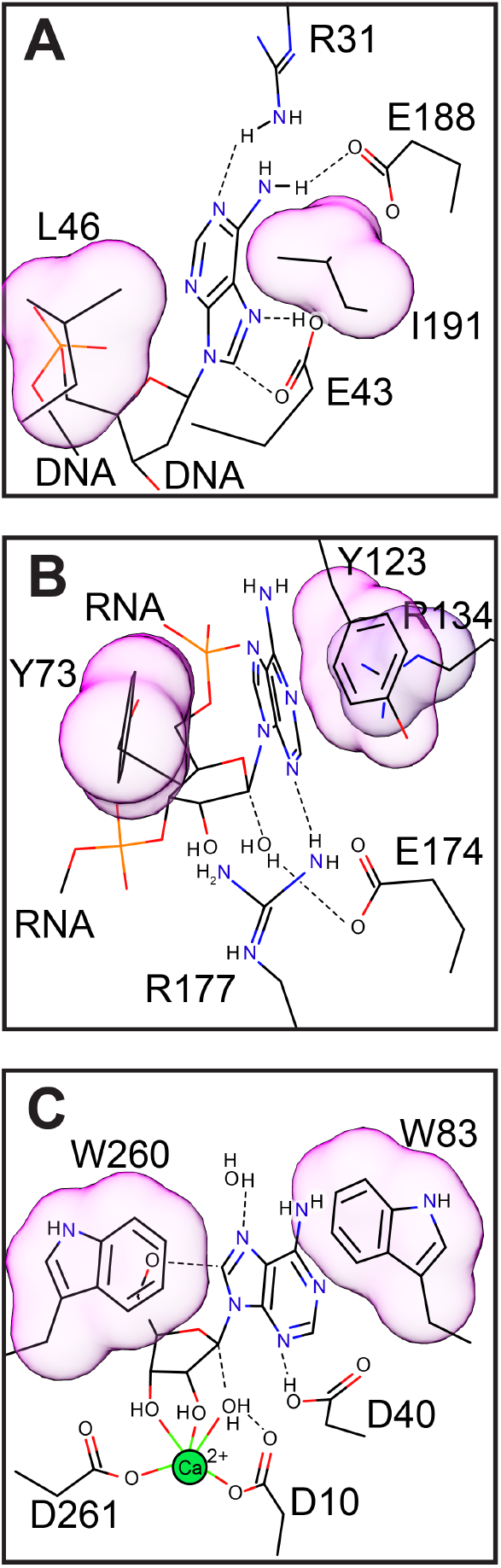
Comparison of DNA, RNA, and nucleoside adenine glycosylases. Adenine-sandwiching residues are highlighted. **A.** MutY excises adenine from DNA and sandwiches the nucleobase with non-aromatic, aliphatic residues. **B.** Saporin-L3 excises adenine from RNA and sandwiches the leaving group with two aromatic Tyr residues, supported by an Arg. **C.** Purine specific nucleoside hydrolase sandwiches the nucleobase leaving group with two aromatic Trp residues. Drawn on the basis of PDB IDs 3g0q,^[19]^ 2ff2,^[46]^ and 3hiw.^[47]^

One residue serves multiple catalytic functions for the MutY mechanism. The catalytic Glu (Glu43) is the acid/base catalyst for activation of the leaving group, activation of the nucleophile, and contributes, along with the catalytic Asp (Asp144), to electrostatic stabilization of the positively charged transition states encountered before and after the covalent intermediate (**Figure 1**). In other words, Glu43 is involved in all of the catalytic tasks. For RIPs and purine-specific NH, these catalytic tasks are delegated to several residues. For example, the RIP saporin-L3 accomplishes leaving group stabilization by an extended *π*-bond stacking interaction involving an Arg and two Tyr residues coupled with protonation at N^3^ by a conserved and catalytically critical Arg.^[46]^ These residues devoted to leaving group activation are different from those that provide electrostatic stabilization of the oxocarbenium ion transition state and nucleophile activation, tasks fulfilled by a catalytic Glu.^[48]^ Comparable to RIPs, the purine-specific NH also stabilizes the leaving group via *π*-bond stacking interactions with two Trp residues,^[47,49]^ which are uninvolved with nucleophile activation or TS stabilization.

The stacking interactions featured in leaving group activation for RIPs and NH are absent for MutY (**Figure 5**). Residues sandwiching the adenine nucleobase in MutY are hydrophobic, aliphatic residues (Leu46 and Ile191 in *Gs* MutY), and aromatic residues are avoided at these positions among bacterial orthologs (**Figure S5**). Inspection of amino acid sequence alignments shows Phe is accepted at these adenine-sanwiching positions at a frequency of ∼1% (51/6143 at position 46; 65/6143 at position 191), indicating aromatic residues fit into this space, but there are no examples with both positions occupied by an aromatic residue. Avoidance of aromatic residues, that are otherwise commonly observed for other adenine glycosylases, may prevent premature, inappropriate adenine excision.

MutY’s retaining mechanism relies on Glu43 for both leaving group and nucleophile activation and thereby necessitates a covalent intermediate, an intermediate that is avoided by the other inverting glycosylases. The different catalytic strategies may reflect different biological constraints guiding evolution. Distributing catalytic tasks among many structural features likely contributed to efficient turnover realized for RIPs, which evolved to destroy structured rRNA and thereby disable numerous ribosomes in a cell as quickly as possible. The biological purpose of nucleotide hydrolyases varies, but a high turnover was apparently desirable. Aromatic residues sandwiching the nucleobase in these adenine glycosylases contribute to catalytic efficiency through stabilizing cation-*π* interactions, in addition to, or in place of acid/base catalysis for leaving group activation.^[49]^

By comparison to the RNA and nucleoside adenine glycosylases, MutY is marked by slow turnover, a result of strong product inhibition.^[27]^ High catalytic turnover was not a priority for MutY as there are few authentic substrates in the cell. Instead, evolution invested in making MutY’s search for rare adenine substrates paired with OG rapid,^[50]^ and avoiding chemically identical but inappropriate adenines. The proposed “on hold” state would provide added layers of protection to ensure MutY serves a DNA-repair function even as it breaks the N-glycosidic bond at chemically correct but informationally misplaced adenines. We further suggest that establishing and releasing such a checkpoint is facilitated by strong reliance on one catalytic residue for leaving group activation. This model predicts that faster, more promiscuous versions of a MutY-like enzyme would be created by replacing both of the adenine-sandwiching residues with aromatic residues.

## Conclusion

The outcomes described here are summarized in **Figure 6**. Replacement of the catalytic Glu with Ser or Gln reduced the adenine excision rate by three orders of magnitude. Structures of these Glu43 replacement variants in complex with DNA showed the substrate nucleobase in an unfamiliar *anti* conformation and the enzyme-generated AP product in a closed-ring, *alpha* anomer configuration, very different from the *beta* stereoisomer seen for the N146S variant enzyme in crystal structures,^[25]^ and as inferred for the wildtype enzyme on the basis of enzyme-catalyzed methanolysis.^[23]^ We knew from previous work that Glu is critical for facilitating initial steps involving acid/base catalysis to activate the leaving group.^[18,20]^ The results described here are consistent with the same catalytic Glu also activating the nucleophile to attack from the *beta* face, the same face as leaving group departure. Other adenine excising enzymes acting on RNA and nucleosides distribute catalytic tasks for activating the leaving group and activating the nucleophile among many residues to establish an SN1 inverting mechanism, while MutY relies heavily on the one catalytic Glu for both leaving group and nucleophile activation in an SN1 retaining mechanism. The unique features of MutY’s mechanism may be important to establish the pre-licensing “on hold” state that is released upon integrating information from the partner DNA strand, so as to avoid genome instability caused by inappropriate activity on abundant adenines that should be ignored, and allow for licensed DNA restoring function of MutY at rare substrate adenines paired with oxidative lesions.

**Figure 6.**
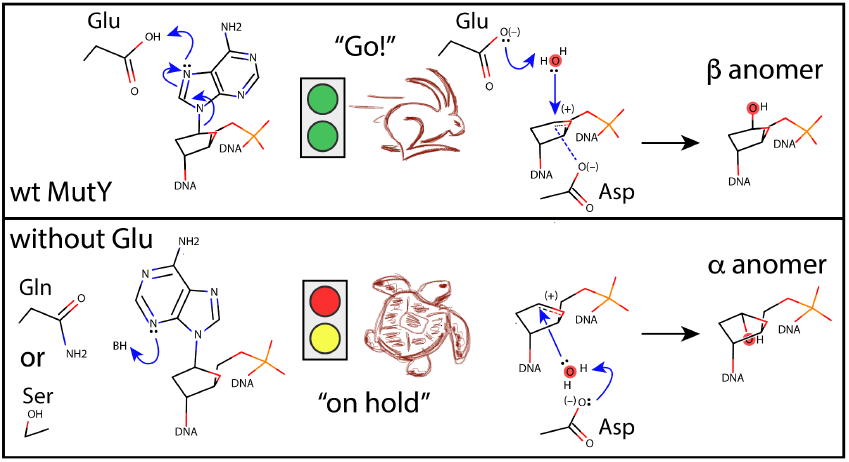
The catalytic Glu activates both the leaving group in a *syn* conformation and the nucleophile for efficient adenine excision with retention of stereochemistry (top). As revealed in this article describing structures and kinetics for E43S and E43Q replacement variants of *Gs* MutY, in the absence of Glu the nucleobase adopts an *anti* conformation, the reaction is very slow, and the resulting AP product is observed in a furanose ring-closed form with *alpha* stereoconfiguration. These features contribute to our understanding of MutY’s function to restore DNA in the face of oxidative damage.

## Experimental Section

See Supplementary Information for the Experimental Section.

## Data and Reagent Availability

Plasmid DNAs have been archived with AddGene. Protein structures have been deposited with the Protein Data Bank.^[51,52]^

## Acknowledgements

Oligonucleotides were synthesized by the DNA/Peptide Facility, part of the Health Sciences Center Cores at the University of Utah. Work was funded by the NSF with awards to SSD CHE-2204228 and MPH CHE-2204229. We thank George Miegs at beamline 8.3.1 and Marc Allaire and Corie Ralston at beamline 5.0.2 for helping with data collection. Beamline 8.3.1 at the Advanced Light Source is operated by the University of California Office of the President, Multicampus Research Programs and Initiatives grant MR-15-328599 the National Institutes of Health (R01 GM124149 and P30 GM124169), Plexxikon Inc. and the Integrated Diffraction Analysis Technologies program of the US Department of Energy Office of Biological and Environmental Research. The Advanced Light Source (Berkeley, CA) is a national user facility operated by Lawrence Berkeley National Laboratory on behalf of the US Department of Energy under contract number DE-AC02-05CH11231, Office of Basic Energy Sciences. The Berkeley Center for Structural Biology is supported in part by the Howard Hughes Medical Institute. The Pilatus detector on 5.0.2 was funded under NIH grant S10OD021832.

## Supplementary Information

The authors have cited additional references within the Supplementary Information.^[14,19,23,25,36,53–63]^

## Footnotes

1 https://www.ncbi.nlm.nih.gov/clinvar/RCV001025999/ https://www.ncbi.nlm.nih.gov/clinvar/RCV000566693/

## References

[1] M. Cabrera, Y. Nghiem, J. H. Miller, “mutM, a second mutator locus in Escherichia coli that generates G.C T.A transversions” J. Bacteriol. 1988, 170, 5405–5407.

[2] S. Boiteux, T. R. O’Connor, J. Laval, “Formamidopyrimidine-DNA glycosylase of Escherichia coli: cloning and sequencing of the fpg structural gene and overproduction of the protein” EMBO J. 1987, 6, 3177–3183.

[3] S. D. Bruner, D. P. Norman, G. L. Verdine, “Structural basis for recognition and repair of the endogenous mutagen 8-oxoguanine in DNA” Nature 2000, 403, 859–866.

[4] F. Faucher, S. Duclos, V. Bandaru, S. S. Wallace, S. Doublié, “Crystal structures of two archaeal 8-oxoguanine DNA glycosylases of the Ogg2 family provide structural insight into guanine/8-oxoguanine distinction” Struct. Lond. Engl. 1993 2009, 17, 703–712.

[5] F. Faucher, S. Doublié, Z. Jia, “8-oxoguanine DNA glycosylases: one lesion, three subfamilies” Int. J. Mol. Sci. 2012, 13, 6711–6729.

[6] H. Maki, M. Sekiguchi, “MutT protein specifically hydrolyses a potent mutagenic substrate for DNA synthesis” Nature 1992, 355, 273–275.

[7] R. G. Fowler, R. M. Schaaper, “The role of the mutT gene of Escherichia coli in maintaining replication fidelity” FEMS Microbiol. Rev. 1997, 21, 43–54.

[8] D. M. Banda, N. N. Nuñez, M. A. Burnside, K. M. Bradshaw, S. S. David, “Repair of 8-oxoG:A Mismatches by the MUTYH Glycosylase: Mechanism, Metals & Medicine” Free Radic. Biol. Med. 2017, 107, 202–215.

[9] M. L. Michaels, C. Cruz, A. P. Grollman, J. H. Miller, “Evidence that MutY and MutM combine to prevent mutations by an oxidatively damaged form of guanine in DNA” Proc. Natl. Acad. Sci. U. S. A. 1992, 89, 7022–7025.

[10] A. L. Livingston, V. L. O’Shea, T. Kim, E. T. Kool, S. S. David, “Unnatural substrates reveal the importance of 8-oxoguanine for in vivo mismatch repair by MutY” Nat. Chem. Biol. 2008, 4, 51–58.

[11] S. G. Conlon, C. Khuu, C. H. Trasviña-Arenas, T. Xia, M. L. Hamm, A. G. Raetz, S. S. David, “Cellular Repair of Synthetic Analogs of Oxidative DNA Damage Reveals a Key Structure–Activity Relationship of the Cancer-Associated MUTYH DNA Repair Glycosylase” ACS Cent. Sci. 2024, 10, 291–301.

[12] N. Al-Tassan, N. H. Chmiel, J. Maynard, N. Fleming, A. L. Livingston, G. T. Williams, A. K. Hodges, D. R. Davies, S. S. David, J. R. Sampson, J. P. Cheadle, “Inherited variants of MYH associated with somatic G:C→T:A mutations in colorectal tumors” Nat. Genet. 2002, 30, 227–232.

[13] A. G. Raetz, S. S. David, “When you’re strange: Unusual features of the MUTYH glycosylase and implications in cancer” DNA Repair 2019, 80, 16–25.

[14] L. P. Russelburg, V. L. O’Shea Murray, M. Demir, K. R. Knutsen, S. L. Sehgal, S. Cao, S. S. David, M. P. Horvath, “Structural Basis for Finding OG Lesions and Avoiding Undamaged G by the DNA Glycosylase MutY” ACS Chem. Biol. 2020, 15, 93–102.

[15] P. H. Utzman, V. P. Mays, B. C. Miller, M. C. Fairbanks, W. J. Brazelton, M. P. Horvath, “Metagenome mining and functional analysis reveal oxidized guanine DNA repair at the Lost City Hydrothermal Field” PLOS ONE 2024, 19, e0284642.

[16] C. H. Trasviña-Arenas, M. Demir, W.-J. Lin, S. S. David, “Structure, function and evolution of the Helix-hairpin-Helix DNA glycosylase superfamily: Piecing together the evolutionary puzzle of DNA base damage repair mechanisms” DNA Repair 2021, 108, 103231.

[17] C. Majumdar, M. Demir, S. R. Merrill, M. Hashemian, S. S. David, “FSHing for DNA Damage: Key Features of MutY Detection of 8-Oxoguanine:Adenine Mismatches” Acc. Chem. Res. 2024, 57, 1019–1031.

[18] J. A. B. McCann, P. J. Berti, “Transition-state analysis of the DNA repair enzyme MutY” J. Am. Chem. Soc. 2008, 130, 5789–5797.

[19] S. Lee, G. L. Verdine, “Atomic substitution reveals the structural basis for substrate adenine recognition and removal by adenine DNA glycosylase” Proc. Natl. Acad. Sci. 2009, 106, 18497–18502.

[20] M. K. Brinkmeyer, M. A. Pope, S. S. David, “Catalytic Contributions of Key Residues in the Adenine Glycosylase MutY Revealed by pH-dependent Kinetics and Cellular Repair Assays” Chem. Biol. 2012, 19, 276–286.

[21] C. Majumdar, N. N. Nuñez, A. G. Raetz, C. Khuu, S. S. David, “Cellular assays for studying the Fe-S cluster containing base excision repair glycosylase MUTYH and homologs” Methods Enzymol. 2018, 599, 69–99.

[22] M. J. Landrum, J. M. Lee, G. R. Riley, W. Jang, W. S. Rubinstein, D. M. Church, D. R. Maglott, “ClinVar: public archive of relationships among sequence variation and human phenotype” Nucleic Acids Res. 2014, 42, D980–985.

[23] R. D. Woods, V. L. O’Shea, A. Chu, S. Cao, J. L. Richards, M. P. Horvath, S. S. David, “Structure and stereochemistry of the base excision repair glycosylase MutY reveal a mechanism similar to retaining glycosidases” Nucleic Acids Res. 2016, 44, 801–810.

[24] M. D. Hanwell, D. E. Curtis, D. C. Lonie, T. Vandermeersch, E. Zurek, G. R. Hutchison, “Avogadro: an advanced semantic chemical editor, visualization, and analysis platform” J. Cheminformatics 2012, 4, 17.

[25] M. Demir, L. P. Russelburg, W.-J. Lin, C. H. Trasviña-Arenas, B. Huang, P. K. Yuen, M. P. Horvath, S. S. David, “Structural snapshots of base excision by the cancer-associated variant MutY N146S reveal a retaining mechanism” Nucleic Acids Res. 2023, 51, 1034–1049.

[26] D. J. Nikkel, S. D. Wetmore, “Distinctive Formation of a DNA-Protein Cross-Link during the Repair of DNA Oxidative Damage: Insights into Human Disease from MD Simulations and QM/MM Calculations” J. Am. Chem. Soc. 2023, 145, 13114–13125.

[27] S. L. Porello, A. E. Leyes, S. S. David, “Single-turnover and pre-steady-state kinetics of the reaction of the adenine glycosylase MutY with mismatch-containing DNA substrates” Biochemistry 1998, 37, 14756–14764.

[28] A. R. Fersht, “Dissection of the structure and activity of the tyrosyl-tRNA synthetase by site-directed mutagenesis” Biochemistry 1987, 26, 8031–8037.

[29] J. R. Knowles, “Tinkering with enzymes: what are we learning?” Science 1987, 236, 1252–1258.

[30] B. V. Plapp, “Site-directed mutagenesis: a tool for studying enzyme catalysis” Methods Enzymol. 1995, 249, 91–119.

[31] A. Peracchi, “Enzyme catalysis: removing chemically ‘essential’ residues by site-directed mutagenesis” Trends Biochem. Sci. 2001, 26, 497–503.

[32] P. Carter, J. A. Wells, “Dissecting the catalytic triad of a serine protease” Nature 1988, 332, 564–568.

[33] L. Wang, S.-J. Lee, G. L. Verdine, “Structural Basis for Avoidance of Promutagenic DNA Repair by MutY Adenine DNA Glycosylase” J. Biol. Chem. 2015, 290, 17096–17105.

[34] C. H. Trasviña-Arenas, U. C. Dissanayake, N. Tamayo, M. Hashemian, W. J. Lin, M. Demir, N. Hoyos-Gonzalez, A. J. Fisher, G. A. Cisneros, M. P. Horvath, S. S. David, “Structure of human MUTYH and functional profiling of cancer-associated variants reveal an allosteric network between its [4Fe-4S] cluster cofactor and active site required for DNA repair” Nat. Commun. 2025, 16, 3596.

[35] J. C. Fromme, A. Banerjee, S. J. Huang, G. L. Verdine, “Structural basis for removal of adenine mispaired with 8-oxoguanine by MutY adenine DNA glycosylase” Nature 2004, 427, 652–656.

[36] A. W. Francis, S. A. Helquist, E. T. Kool, S. S. David, “Probing the requirements for recognition and catalysis in Fpg and MutY with nonpolar adenine isosteres” J. Am. Chem. Soc. 2003, 125, 16235–16242.

[37] K. G. Berdal, R. F. Johansen, E. Seeberg, “Release of normal bases from intact DNA by a native DNA repair enzyme” EMBO J. 1998, 17, 363–367.

[38] M. D. Wyatt, J. M. Allan, A. Y. Lau, T. E. Ellenberger, L. D. Samson, “3-methyladenine DNA glycosylases: structure, function, and biological importance” BioEssays News Rev. Mol. Cell. Dev. Biol. 1999, 21, 668–676.

[39] S. Ej, P. Jl, W. Sd, “Effects of nucleophile, oxidative damage, and nucleobase orientation on the glycosidic bond cleavage in deoxyguanosine” J. Phys. Chem. B 2010, 114, DOI 10.1021/jp9113656.

[40] S. A. P. Lenz, J. L. Kellie, S. D. Wetmore, “Glycosidic Bond Cleavage in DNA Nucleosides: Effect of Nucleobase Damage and Activation on the Mechanism and Barrier” J. Phys. Chem. B 2015, 119, 15601–15612.

[41] D. J. Vocadlo, G. J. Davies, R. Laine, S. G. Withers, “Catalysis by hen egg-white lysozyme proceeds via a covalent intermediate” Nature 2001, 412, 835–838.

[42] N. J. Rzechorzek, S. Kunzelmann, A. G. Purkiss, M. Silva Dos Santos, J. I. MacRae, I. A. Taylor, K. Fugger, S. C. West, “Mechanism of substrate hydrolysis by the human nucleotide pool sanitiser DNPH1” Nat. Commun. 2023, 14, 6809.

[43] H. Yuan, C. F. Stratton, V. L. Schramm, “Transition State Structure of RNA Depurination by Saporin L3” ACS Chem. Biol. 2016, 11, 1383–1390.

[44] X. Y. Chen, P. J. Berti, V. L. Schramm, “Ricin A-chain: Kinetic isotope effects and transition state structure with stem-loop RNA” J. Am. Chem. Soc. 2000, 122, 1609–1617.

[45] P. J. Berti, J. A. B. McCann, “Toward a detailed understanding of base excision repair enzymes: transition state and mechanistic analyses of N-glycoside hydrolysis and N-glycoside transfer” Chem. Rev. 2006, 106, 506–555.

[46] M.-C. Ho, M. B. Sturm, S. C. Almo, V. L. Schramm, “Transition state analogues in structures of ricin and saporin ribosome-inactivating proteins” Proc. Natl. Acad. Sci. 2009, 106, 20276–20281.

[47] W. Versées, J. Barlow, J. Steyaert, “Transition-state complex of the purine-specific nucleoside hydrolase of T. vivax: enzyme conformational changes and implications for catalysis” J. Mol. Biol. 2006, 359, 331–346.

[48] A. Frankel, P. Welsh, J. Richardson, J. D. Robertus, “Role of arginine 180 and glutamic acid 177 of ricin toxin A chain in enzymatic inactivation of ribosomes” Mol. Cell. Biol. 1990, 10, 6257–6263.

[49] W. Versées, S. Loverix, A. Vandemeulebroucke, P. Geerlings, J. Steyaert, “Leaving Group Activation by Aromatic Stacking: An Alternative to General Acid Catalysis” J. Mol. Biol. 2004, 338, 1–6.

[50] A. J. Lee, C. Majumdar, S. D. Kathe, R. P. Van Ostrand, H. R. Vickery, A. M. Averill, S. R. Nelson, A. H. Manlove, M. A. McCord, S. S. David, “Detection of OG:A Lesion Mispairs by MutY Relies on a Single His Residue and the 2-Amino Group of 8-Oxoguanine” J. Am. Chem. Soc. 2020, 142, 13283–13287.

[51] H. M. Berman, J. Westbrook, Z. Feng, G. Gilliland, T. N. Bhat, H. Weissig, I. N. Shindyalov, P. E. Bourne, “The Protein Data Bank” Nucleic Acids Res. 2000, 28, 235–242.

[52] H. Berman, K. Henrick, H. Nakamura, “Announcing the worldwide Protein Data Bank” Nat. Struct. Mol. Biol. 2003, 10, 980–980.

[53] W. Kabsch, “XDS” Acta Crystallogr. D Biol. Crystallogr. 2010, 66, 125–132.

[54] W. Kabsch, “Integration, scaling, space-group assignment and post-refinement” Acta Crystallogr. D Biol. Crystallogr. 2010, 66, 133–144.

[55] D. Liebschner, P. V. Afonine, M. L. Baker, G. Bunkóczi, V. B. Chen, T. I. Croll, B. Hintze, L.-W. Hung, S. Jain, A. J. McCoy, N. W. Moriarty, R. D. Oeffner, B. K. Poon, M. G. Prisant, R. J. Read, J. S. Richardson, D. C. Richardson, M. D. Sammito, O. V. Sobolev, D. H. Stockwell, T. C. Terwilliger, A. G. Urzhumtsev, L. L. Videau, C. J. Williams, P. D. Adams, “Macromolecular structure determination using X-rays, neutrons and electrons: recent developments in Phenix” Acta Crystallogr. Sect. Struct. Biol. 2019, 75, 861–877.

[56] P. Emsley, B. Lohkamp, W. G. Scott, K. Cowtan, “Features and development of Coot” Acta Crystallogr. D Biol. Crystallogr. 2010, 66, 486–501.

[57] N. W. Moriarty, E. J. Draizen, P. D. Adams, “An editor for the generation and customization of geometry restraints” Acta Crystallogr. Sect. Struct. Biol. 2017, 73, 123–130.

[58] K. Diederichs, P. A. Karplus, “Improved R-factors for diffraction data analysis in macromolecular crystallography” Nat. Struct. Biol. 1997, 4, 269–275.

[59] L. Li, J. W. Szostak, “The free energy landscape of pseudorotation in 3’-5’ and 2’-5’ linked nucleic acids” J. Am. Chem. Soc. 2014, 136, 2858–2865.

[60] A. Z. Michelson, A. Rozenberg, Y. Tian, X. Sun, J. Davis, A. W. Francis, V. L. O’Shea, M. Halasyam, A. H. Manlove, S. S. David, J. K. Lee, “Gas-phase studies of substrates for the DNA mismatch repair enzyme MutY” J. Am. Chem. Soc. 2012, 134, 19839–19850.

[61] C. Majumdar, P. L. McKibbin, A. E. Krajewski, A. H. Manlove, J. K. Lee, S. S. David, “Unique Hydrogen Bonding of Adenine with the Oxidatively Damaged Base 8-Oxoguanine Enables Specific Recognition and Repair by DNA Glycosylase MutY” J. Am. Chem. Soc. 2020, 142, 20340–20350.

[62] F. Madeira, N. Madhusoodanan, J. Lee, A. Eusebi, A. Niewielska, A. R. N. Tivey, R. Lopez, S. Butcher, “The EMBL-EBI Job Dispatcher sequence analysis tools framework in 2024” Nucleic Acids Res. 2024, 52, W521–W525.

[63] G. E. Crooks, G. Hon, J.-M. Chandonia, S. E. Brenner, “WebLogo: a sequence logo generator” Genome Res. 2004, 14, 1188–1190.

