## Supporting Information for "Structural Basis for Nucleobase Activation by the Adenine DNA Glycosylase MutY"

**Table S1.** Data collection and model refinement statistics

| Data collection statistic | S43:Adenine | Q43:Purine | S43:Abasic |
|---|---|---|---|
| Wavelength (Å) | 1.116 | 1.116 | 1.000 |
| Resolution range (Å) | 50.14 - 2.6 | 68.98 - 2.2 | 42.96 - 1.68 |
| (Highest resolution) | (2.693 - 2.6) | (2.279 - 2.2) | (1.74 - 1.68) |
| Space group | P 1 21 1 | P 1 21 1 | P 21 21 21 |
| Unit cell a, b, c (Å) | 49.08 137.9 74.52 | 48.63 137.96 74.49 | 37.74 85.92 140.69 |
| *beta* (°) | 101.436 | 101.000 | 90 |
| Total reflections | 150327 (10956) | 328794 (24485) | 679161 (51832) |
| Unique reflections (+) (-) | 58172 (4259) | 96293 (7112) | 100381 (7356) |
| Unique reflections | 29767 (2942) | 48807 (4834) | 53020 (5165) |
| Multiplicity | 2.6 (2.6) | 3.4 (3.4) | 6.8 (7.0) |
| Completeness (%) | 99.68 (99.12) | 99.92 (99.96) | 99.64 (98.81) |
| Mean I/sigma(I) | 5.6 (0.8) | 5.9 (0.9) | 9.4 (1.1) |
| Wilson B-factor ($Å^2$) | 55.18 | 48.18 | 30.73 |
| R-merge (%) | 19.1 (207) | 11.5 (142) | 9.0 (191) |
| R-meas (%) ‡ | 24.0 (262) | 13.6 (168) | 9.8 (205) |
| CC1/2 (%) | 98.1 (9.1) | 99.3 (30.0) | 99.8 (31.3) |
| **Model refinement statistic** | **S43:Adenine** | **Q43:Purine** | **S43:Abasic** |
| Reflections used in refinement | 29763 (2942) | 48787 (4834) | 53019 (5165) |
| Reflections used for R-free | 771 (79) | 1234 (123) | 1328 (130) |
| R-work | 0.2354 (0.3499) | 0.2080 (0.3392) | 0.2207 (0.3742) |
| R-free | 0.2607 (0.3496) | 0.2400 (0.3707) | 0.2483 (0.3879) |
| Number of non-hydrogen atoms | 6373 | 6509 | 3344 |
| MutY | 5374 | 5427 | 2681 |
| DNA | 879 | 861 | 422 |
| 8OG | 46 | 46 | 23 |
| Active site DNA | 42 | 40 | 12 |

| **Table S1.** Data collection and model refinement statistic (continued) | | | |
|---|---|---|---|
| **Model refinement statistic** | **S43:Adenine** | **Q43:Purine** | **S43:Abasic** |
| Number of non-hydrogen atoms calcium | 2 | 5 | 2 |
| 4Fe4S | 16 | 16 | 16 [€] |
| solvent | 98 | 200 | 224 |
| Protein residues | 701 | 702 | 346 |
| RMS(bonds) | 0.003 | 0.003 | 0.01 |
| RMS(angles) | 0.53 | 0.66 | 1.24 |
| Ramachandran favored (%) | 97.4 | 97.1 | 96.5 |
| Ramachandran allowed (%) | 2.6 | 2.9 | 3.5 |
| Ramachandran outliers (%) | 0 | 0 | 0 |
| Rotamer outliers (%) | 0.2 | 0 | 1.1 |
| Clashscore | 1.2 | 0.9 | 6.2 |
| Number of TLS groups | 4 | 4 | 3 |
| Average B-factor ($Å^2$) | 57.7 | 58.2 | 49.6 |
| MutY | 59.5 | 60.0 | 51.1 |
| DNA | 48.0 | 48.9 | 43.3 |
| 8OG | 39.2 | 40.7 | 26.6 |
| active site DNA | 52.0 | 60.3 | 33.1 |
| calcium ion | 50.5 | 60.0 | 63.5 |
| 4FeS4 | 52.8 | 51.1 | 38.0 |
| solvent | 50.2 | 50.5 | 45.2 |
| Occupancy of active site nucleobase | 1.00, 1.00 | 0.84, 0.73 | 0.00 |

[‡] Redundancy independent measure of R.[58]
[€] 4Fe-4S modeled with two alternate conformations.

**Table S2**. Glycosylase activity detected by X-ray crystallography

| Variant | Starting DNA | Evaluation of DNA in active site | Number of crystals |
|---|---|---|---|
| E43S | OG:A | Substrate, *anti* conformer | 4 |
| E43S | OG:P | AP Product, *alpha* anomer | 7 |
| E43Q | OG:A | NA [a] | 0 |
| E43Q | OG:P | Substrate, partial occupancy, *anti* conformer | 3 |

[a] E43Q complexed with OG:dA DNA failed to crystallize

**Glycosylase activity for E43S and E43Q *Gs* MutY acting on OG:dP**

With the purine substrate these enzymes are even slower, in fact so impaired that instead of precise rate determination we left reactions to proceed for a full day and compared rates by comparing the extent of observed product accumulation after 3h, 6h, and 24h. We observed three to nine-fold greater product accumulation for the enzyme catalyzed reactions in comparison to the no enzyme control. E43Q produced 19% cleaved product and E43S produced 54% cleaved product after a 24 hour period compared to the no enzyme control of 6% cleaved product (**Figure S1**). After subtracting background, these values yielded estimates of 0.0002 $min^{-1}$ and 0.001 $min^{-1}$ for the upper limit of purine excision for the E43Q and E43S enzymes, respectively. The E43S enzyme tolerated the switch to a purine substrate better than E43Q, although both were severely impacted. In summary, replacement of Glu43 makes the enzyme extremely slow, especially with the purine substrate, yet the variant *Gs* MutY enzymes retain residual catalytic activity contrary to previous work with the *E. coli* enzyme.

| | **E43Q *Gs* MutY** | | | **E43S *Gs* MutY** | | | **No Enzyme** | | |
|---|---|---|---|---|---|---|---|---|---|
| Time | 3 hr | 6 hr | 24 hr | 3 hr | 6 hr | 24 hr | 3 hr | 6 hr | 24 hr |
| Percent Cleavage | 6% | 9% | 19% | 17% | 32% | 54% | 6.4% | 3.5% | 6.2% |

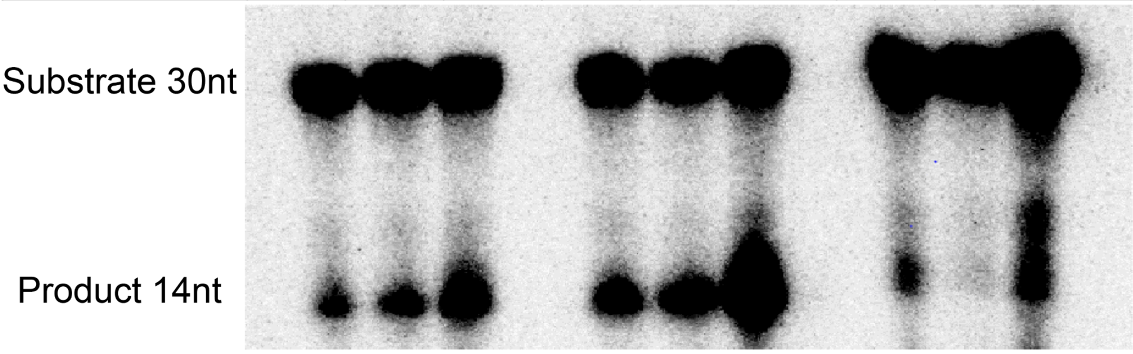

**Figure S1**. Glycosylase reactions with E43Q and E43S processing the OG:dP substrate. Radiolabelled dsDNA containing OG across from purine was combined with enzyme, incubated at 60 °C, and reactions were quenched by addition of sodium hydroxide to cleave DNA at the AP product. Bands correspond to the low-mobility substrate DNA (30 nt) and high-mobility product DNA (14 nt). Percent cleavage values were calculated after subtracting the no-enzyme control. The E43S and E43Q variants show glycosylase activity that is impaired relative to the wildtype enzyme but not completely dead.

#### Distinguishing *anti/syn* conformations for substrate nucleobase

Discovery maps calculated early in the structure refinement process for $E43S_{OG:dA}$ reveal the outline of an adenine base in an *anti*-like conformation with the six-member ring pointed away from the sugar moiety (**Figure S2A**). A starting model with a *syn* nucleobase would emerge in this *anti* conformation following reciprocal space refinement with *Phenix* (**Figure S2B**). Obtaining a model with the *syn* conformation required increased torsion angle restraints to freeze rotation about the $C^{1'}$-$N^{9}$ glycosidic bond. Difference maps calculated for such a model showed clear negative and positive electron density features (**Figure S2C**) that indicate the *anti* conformation predominates in structures for the Glu replacement variant enzyme.

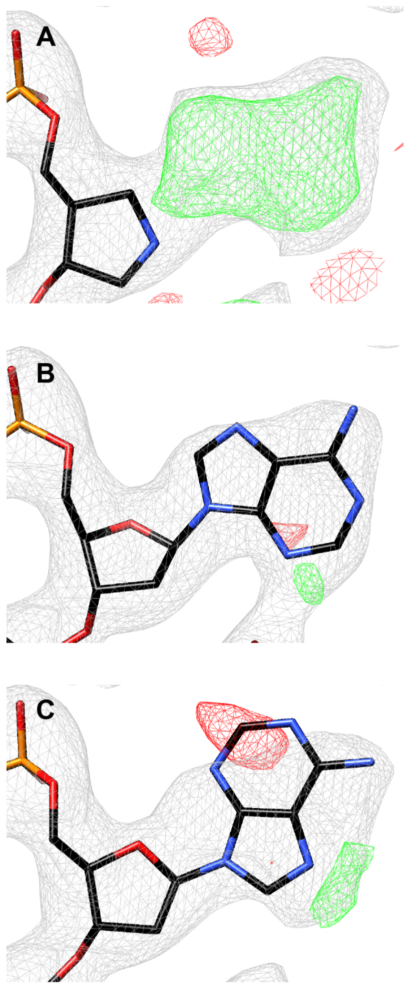

**Figure S2.** Adenine adopts an *anti* orientation **A.** Map calculated at the active site of $E43S_{OG:dA}$ after molecular replacement with *Gs* MutY complexed with DNA containing the abasic transition state mimic 1N (PDB ID 6u7t). **B.** Adenine in its *anti* conformation fits the 2|Fo| - |Fc| map with few features in the. **C.** The *syn* conformation poorly fits the the electron density map, with significant positive and negative peaks in the |Fo| - |Fc| difference map (red, green).

### Alternative mechanisms

The *alpha* anomer generated by the E43S variant enzyme suggests an inverting mechanism may contribute to product accumulation. In the absence of Glu43, an inverting mechanism for E43S may initiate with protonation at $N^3$ of the nucleobase in its *anti* conformation. Prior structure activity relation (SAR) studies with substrate analogs showed the importance of hydrogen bonding with $N^3$ for adenine excision by MutY.[36,60,61] With replacement of Glu43 and adoption of the *anti* conformation with $C^{3'}$-*exo* sugar pucker, the water-mediated hydrogen bond network involving $N^3$ observed for the wildtype enzyme is replaced by a direct hydrogen bond to Tyr126 (**Figure 4**), and we speculate that Tyr126 serves as the proton donor to activate the leaving group (**Figure S3**). The inverting mechanism proceeds with nucleophile attacking from the 3' face, facilitated by Asp144 (**Figure S3**, top branch), as proposed originally for MutY.[19] Alternatively, AP product accumulation in the *alpha* anomer may result from the now established inverting mechanism for MutY,[23] with Tyr126 or some other acid/base catalyst serving to activate the nucleophile (**Figure S3**, bottom branch), followed by racemization *via* the ring-open aldehyde intermediate and selection for the *alpha* anomer by the enzyme. However, this alternate route is made less plausible when the *alpha* anomer AP product observed for E43S is compared to the *beta* anomer AP product observed for N146S (see below, **Figure S4**). As the interactions available for stereospecific selection from a racemic mixture are essentially the same for E43S and N146S, its difficult to explain the distinct stereoisomers observed in electron density maps for these different MutY variants. Therefore, we favor the more direct route to the *alpha* anomer observed for the structure of E43S$_{OG:AP}$ described in this work, but cannot exclude alternate routes.

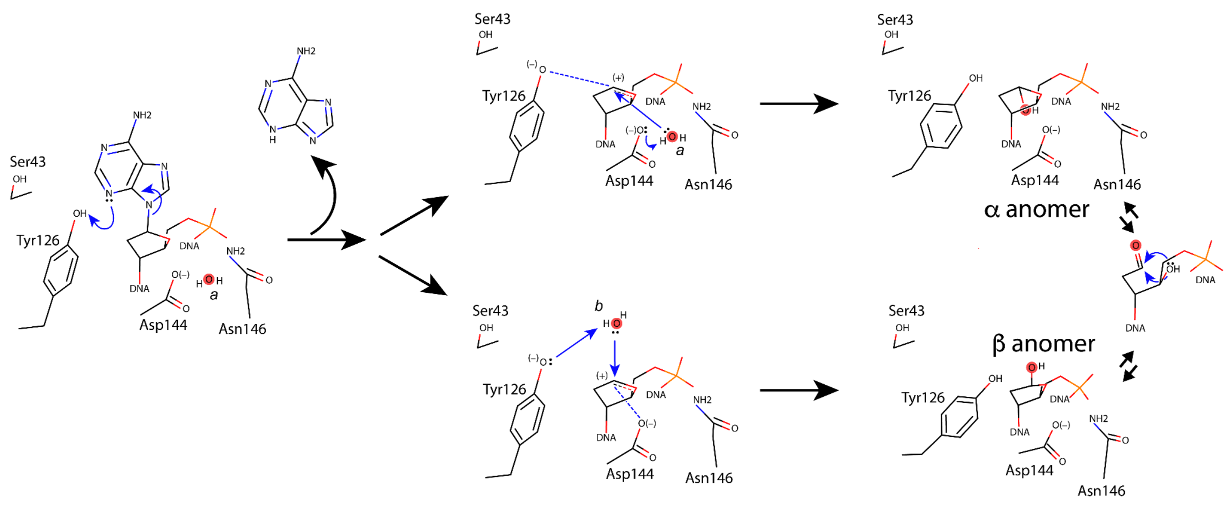

**Figure S3**. Catalytic routes to an *alpha* AP site for E43S. In the absence of a catalytic Glu, an alternate residue (we suggest Tyr126) protonates the nucleobase in its *anti*-like conformation. Two alternate routes diverge following leaving group activation and departure. In the top branch, Asp144 activates the nucleophile (*a*), and Tyr126 in its alkoxide form stabilizes the transition state that resolves with attack from the 3'-face to yield the *alpha* AP site directly. An indirect route follows the bottom branch. In this indirect route, Tyr126 activates the nucleophile (*b*), and Asp144 stabilizes the oxocarbenium ion transition state that resolves with attack from the 5'-face to yield the *beta* anomer, which interconverts *via* the ring-open aldehyde intermediate. In the indirect route, interactions with the enzyme active site selects the *alpha* anomer from the racemic mixture. Of these two possibilities, we favor the direct route because interactions with active site residues are highly comparable for E43S and N146S variants of Gs MutY, yet the stereoconfiguration of the enzyme-generated AP site is *alpha* for E43S and *beta* for N146S (**Figure S4**).

### Comparison of enzyme-generated AP product for N146S and E43S

We recently described structures for *Gs* MutY with the cancer associated N146S amino acid replacement.[25] One of these reveals the enzyme-generated AP product refined with data measured to the 1.68-Å resolution limit, sufficient to distinguish the ring-closed furanose *beta* anomer with a C4'-*exo* sugar pucker (**Figure S4A**). The electron density map for $E46S_{OG:AP}$ (**Figure S4B**), also calculated to the 1.68-Å resolution limit, is very different from that of $N146S_{OG:AP}$ (**Figure S4A**) in the region of the AP site and at the substituting positions. The catalytic Asp (D144) and supporting Tyr (Y126) residues are positioned similarly in the two structures, as defined by clear electron density. These are the only hydrogen-bonding partners available to the hydroxyl group of the AP site in both structures; Glu 43 is too distant to make a significant interaction. Selecting the *alpha* anomer or the *beta* anomer from a racemized mixture seems an unlikely explanation for the different outcomes observed for $N146S_{OG:AP}$ and $E43S_{OG:AP}$, given the very similar positions of Asp144 and Tyr126. It is reasonable and plausible, therefore, to suggest that the AP product seen in electron density maps is the stereoisomer generated by enzyme catalysis.

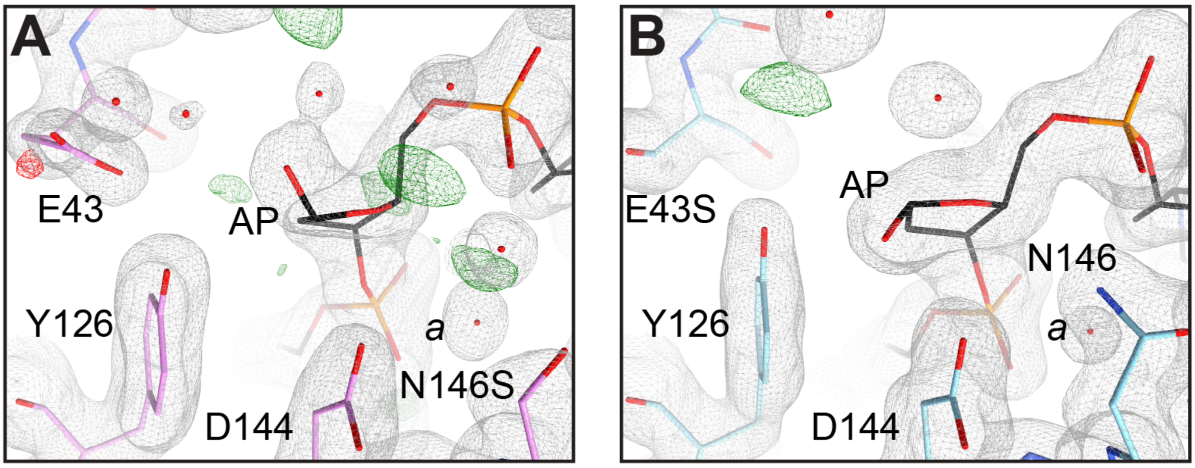

**Figure S4**. Comparison of enzyme-generated AP sites. Electron density maps were calculated to the 1.68 Å resolution limit and contoured at the 1.2-sigma level for 2|Fo| - |Fc| maps (grey) and at the +/- 3.8-sigma level for |Fo|-|Fc| maps (green, red). The N146S-generated AP product adopts the *beta* anomer consistent with retention of stereoconfiguration (A) while the E43S-generated AP product adopts the *alpha* anomer consistent with inversion of stereoconfiguration (B).

### Experimental Section

#### Sequence conservation analysis

The amino acid sequences of MutY from 6143 different bacteria species were harvested from KEGG and split into groups of 200 before alignment with MUSCLE.[62] Each alignment included the *Gs* MutY reference sequence. The two motifs corresponding to positions 41-51 and positions 186-196 in *Gs* MutY were located and analyzed for amino acid type with an *R* program written by the author M.P.H. The LOGOs shown in **Figure S5** were obtained with the WebLogo server,[63] https://weblogo.berkeley.edu/ after randomly selecting 2000 sequences. Inclusion in this analysis required the MutY to be longer than 320 residues and contain Glu or Asp at position 43 (**Figure S5**, E in the first motif). Most of these sequences also retained the iron-sulfur cluster, although this was not a requirement for inclusion. The residues sandwiching the adenine nucleobase were most often Leu at position 46 and most often Met or Ile at position 191 (**Figure S5**, L in the first motif and M/I in the second motif).

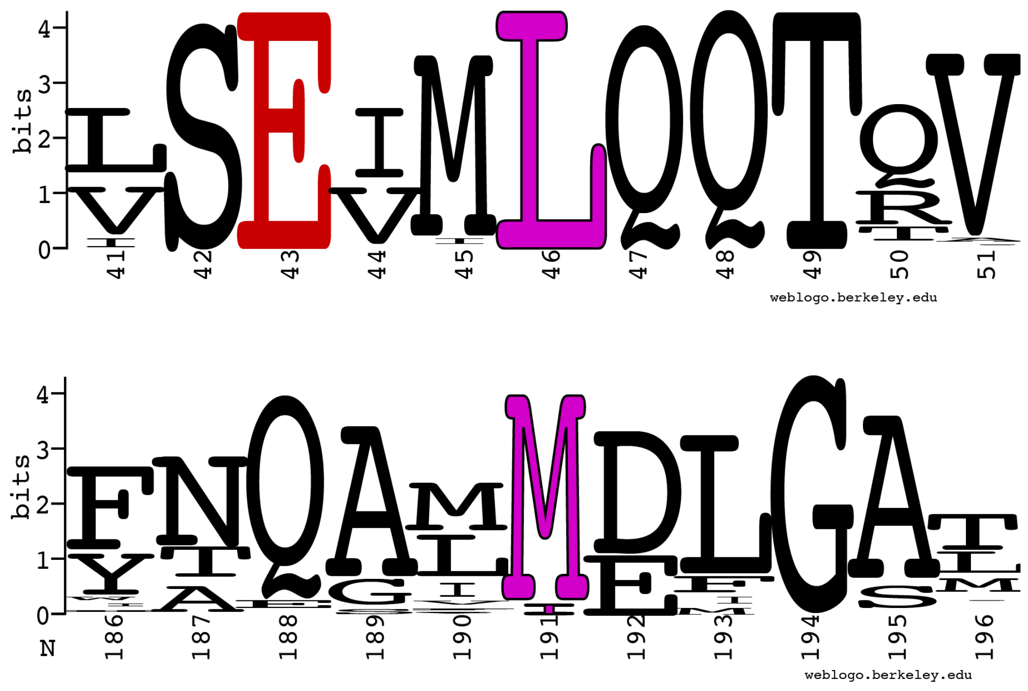

**Figure S5**. Sequence conservation. The catalytic Glu at position 43 is shown in red (top). Residues that sandwich the adenine nucleobase are shown in magenta at position 46 (top) and position 191 (bottom). Out of 6143 MutYs included in this analysis, 51 have an aromatic Phe residue in place of Leu at position 46 and 65 have an aromatic Phe residue in place of M/I at position 191. There were no examples with both positions occupied by an aromatic residue.

**Oligonucleotide synthesis and purification**

8-oxo-dG-CE Phosphoramidite (OG) and 2'-DeoxyNebularine-CE Phosphoramidite (Purine) were purchased from Glen Research. The University of Utah DNA / Peptide core facility synthesized the 11-mer DNAs for crystallography (**Table S3**). Oligonucleotides were decoupled and annealed in 10 mM Tris pH 7.5 as described previously.[14]

**Table S3**. DNA sequences for crystallography

| DNA | Sequence |
|---|---|
| V11OG | 5' - AAGAC(dOG)TGGAC - 3' |
| cV11A | 5' - TGTCCA(dA)GTCT - 3' |
| cV11P | 5' - TGTCCA(dP)GTCT - 3' |

**Molecular cloning**

The pET28 plasmid with T7 promoter driving expression of wildtype *Gs* MutY with an N-terminal 6xHis tag and thrombin cleavage site was generously provided by the Verdine lab.  Site directed mutagenesis to replace the codon for Glu43 with a codon for Ser was completed with the QuickChange kit (QiaGen) according to manufacturer's instructions and with primer DNAs listed in **Table S4**. Ligation independent cloning was applied to generate the pET28 plasmid for E43Q *Gs* MutY expression. Two PCR reactions with primers listed in **Table 4** created overlapping DNAs for the left-hand (LH) and right-hand (RH) halves of the pET28a(+) plasmid. The forward primer for the RH reaction (E43Q-crick) and the reverse primer for the LH reaction (E43Q-watson-franklin) each encoded a Gln codon at position 43 of *Gs* MutY. These primers were paired with oligo DNAs complementary to a sequence immediately downstream of the *ori* origin of replication of the pET28a(+) vector (CW-mph, CCW-mph). PCR products were separated by gel electrophoresis, bands of the correct length were recovered and DNA was purified with the GeneJet kit (Thermo Scientific) according to manufacturer's instructions. The purified PCR products were mixed and directly transformed into competent DH5α cells. Kanamycin resistant transformants were recovered by selection on LB plates with 40 µg / mL kanamycin and the sequence encoding E43Q Gs MutY with N-terminal 6xHis tag was confirmed by Sanger sequencing by the University of Utah sequencing core using the primers T7trm and T7.2_crk (**Table 4**).

**Table S4**. DNA primers for molecular cloning

| **DNA** | **Sequence** |
|---|---|
| E43Q-crick | 5' CCATATAAAGTATGGGTGTCG**C**AAGTGATGCTGCAGCAAACG 3' |
| E43Q-watson-franklin | 5' CGTTTGCTGCAGCATCACTT**G**CGACACCCATACTTTATATGG 3' |
| CW-mph | 5' GCAACGCGGCCTTTTTACGGTTCC 3' |
| CCW-mph | 5' GGAACCGTAAAAAGGCCGCGTTGC 3' |
| T7-v2 | 5' CGTCCGGCGTAGAGGATCG 3' |
| T7-term | 5' GCTAGTTATTGCTCAGCGG 3' |

**Glycosylase Assay**

The rate constants of Gs MutY and its variants were measured by using previously described glycosylase assay methods (ref). A 30 base pair DNA duplex with a central OG:dA or OG:dP mismatch was utilized to determine the rate constants. The 5′-end of the A/P-containing strand was radiolabeled using γ-$^{32}$P-ATP and T4 Polynucleotide Kinase (NEB) and annealed to its complementary strand containing OG in a buffer composition of 20 mM Tris pH 7.6, 10 mM EDTA and 150 mM sodium chloride. The 20 nM duplex was incubated with the enzyme at 60°C in reaction buffer of 20 mM Tris pH 7.6, 10 mM EDTA, 0.1 mg/ml bovine serum albumin (BSA) and 30 mM NaCl. Each aliquot removed from the reaction at certain time points was quenched with sodium hydroxide to introduce a single DNA strand break at the abasic site to form a 14-nucleotide (nt) product strand. Enzyme concentrations of 4, 8 and 12 nM were used for glycosylase assays performed under multiple turnover conditions to measure active enzyme concentration and rate constant of product release, $k_3$. The rate constant of adenine cleavage, $k_2$, was measured under single turnover (STO) conditions where enzyme concentration is higher than DNA concentration. The unprocessed 30-nt substrate strand was separated from the product strand using a denaturing PAGE for quantification of percent product formation.

**X-ray crystallography**

The Glu substitution variants of *Gs* MutY were overexpressed and purified as previously described.[14] *Gs* MutY was exchanged into a buffer containing 20 mM Tris pH 8, 150 mM sodium chloride, and 5 mM *beta*-mercaptoethanol in the final Superdex-200 size-exclusion chromatography. Fractions enriched for MutY as quantified by A280 and A410 were combined with an equal volume of 50% sterile autoclaved glycerol before storing at -80 °C. For crystallization, MutY was concentrated to between 270-360 μM with 10 kDa MWCO ultra centrifugal filters (Millipore). Pure protein was mixed with an equal volume of annealed DNA (525 μM) containing either an OG:dA or OG:Pu lesion. The enzyme-DNA mixture was pre-incubated at various temperatures and times before assembly of crystal trays (**Table S5**). Crystallization well solutions were composed of 100 mM Tris pH 8-9, 12% PEG 4000, 400 mM calcium acetate, 2% ethylene glycol, and 5 mM *beta*-mercaptoethanol. Microseeds were prepared using the Hampton Research Seed Bead kit (HR2-320). Crystals and 50 μL of the well solution were combined in the Seed Bead tube, vortexed for 3 min to completely crush the starting crystals, before increasing the volume by adding 450 μL of a solution matching the original well solution to yield the final 1x microseed stock solution. Microseeds were serially diluted in 10-fold steps with a solution matching the original well solution to yield $10^{-1}$, $10^{-2}$, $10^{-3}$, …, $10^{-5}$ dilutions. Microseeds were either used fresh or the whole dilution series was frozen at -80°C for later use; both methods yielded crystals. We observed some variance in microseed stock performance with some stocks not producing crystals. Microseed stocks that did not produce crystals were discarded and fresh stocks were assembled. We mixed 1μL of the pre-incubated enzyme-DNA mixture with 1μL of microseed dilution to prepare crystallization reactions using the hanging-drop vapor-diffusion method. Yellow to golden crystals, long and flat in shape grew within a few days (**Figure S6**). Crystals were washed briefly with a solution that matched the crystallization well except for the added cryoprotectant, 5% ethylene glycol, before looping and freezing in liquid nitrogen. Crystals were looped with Hampton Research Mounted Cryoloops (0.1-0.2 mm HR4-947) attached to magnetic CrystalCaps (HR4-779) and stored in CrystalCap vials under liquid nitrogen.

**Table S5**. Preincubation and crystallization reaction conditions

| Structure | Preincubation | Well solution pH |
|---|---|---|
| $E43S_{OG:A}$ | 4 °C, 20 h | 8.0 |
| $E43Q_{OG:P}$ | Ambient, 1.5 h | 9.0 |
| $E43S_{OG:AP}$ | Ambient, 0.5 h | 8.5 |

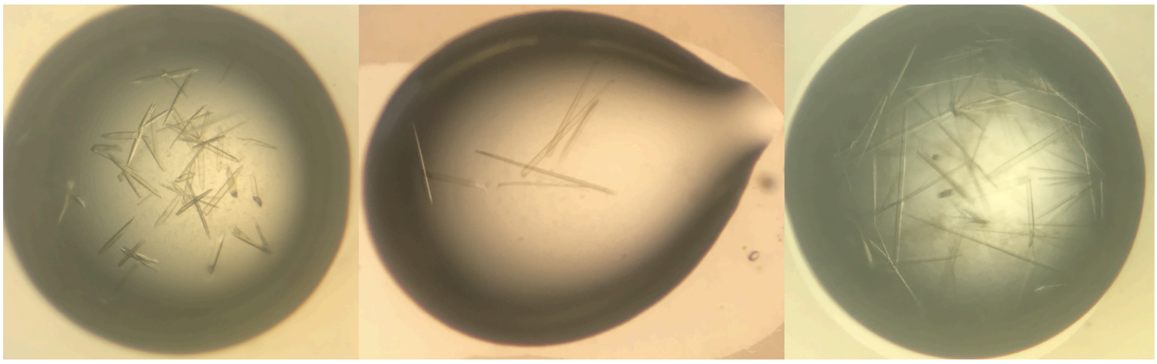

**Figure S6**. Crystals of Glu43 replacement variants of Gs MutY complexed with DNA

Crystals were sent in a dry shipping Dewar to the Advanced Light Source at Berkeley National Laboratory. Data were collected remotely due to the COVID-19 pandemic with Pilatus3 6M detectors at beamlines 5.0.2 and 8.3.1. The highly sensitive next generation detectors enable real time data collection to achieve higher resolution limits. Data were collected in multiple sweeps exposing a crystal at different locations to minimize radiation damage. Most crystals were measured with a strategy of 4 sweeps, each covering 90° in phi, with 0.4 s exposure per frame of 0.2° oscillation. The crystal that yielded the high resolution product structure was measured with a single helical sweep of 360° with 0.25 seconds exposure per frame of 0.25°.

### Data processing

Data integration and processing was accomplished with XDS and XSCALE,[53,54] keeping Bijovet pairs separate so as to calculate anomalous maps useful for distinguishing metals such as calcium ions and the 4Fe-4S metal cluster. For cross-validation, 2.5% of the data were assigned to the test group and these data did not guide minimization of energy functions during model refinement with version 1.20.1 of *Phenix*.[55] Initial phases were obtained by rigid body refinement with a previously refined model. For structures that crystallized in the $P2_1$ space group, the starting model was an unpublished model of MutY-DNA refined in the $P2_1$ space group with crystal id *cg11*. For the product structure which crystallized in the more frequently encountered $P2_12_12_1$ space group, a previously published structure of MutY with PDB ID 6u7t served as the starting model. Both structures feature the transition state analog 1N in the DNA at the position across from OG and thus do not contain a base in the active site. Significantly, this approach minimized concerns with respect to model bias and yielded discovery maps to diagnose presence or absence and pose (if present) of the base. Structure refinement included a round of simulated annealing early in the process to further eliminate model bias. For the high resolution product structure the starting temperature during torsion angle simulated annealing refinement was 900K and the final temperature was 50K, and for the other structures the starting temperature was 1100K and the final temperature 300K. Refinement proceeded with multiple rounds of manually adjusting the models in the context of electron density maps with the model building program *Coot*,[56] followed by positional, temperature factor, and occupancy strategies as implemented with *Phenix Refine*.[55] For the $E43S_{OG:AP}$ structure, the 4Fe-4S metal center and chelating Cys ligands were modeled with two alternate conformations as indicated by anomalous difference maps. For the $E43Q_{OG:dP}$ structure we applied a group occupancy refinement strategy for the purine base so as to refine a common occupancy for all atoms in this group. Parameters defining three TLS groups (DNA, MutY N-terminal domain, MutY C-terminal domain) accounted for anisotropic atom displacement in the final cycle of refinement. Simulated annealing composite omit maps were calculated by the *Composite Omit Map* module of *Phenix*, with 3% of the structure omitted, submitting the remaining structure to torsion angle simulated annealing starting from a temperature of 1100 K, calculating the 2|Fo| - |Fc| map for the omitted region, and repeating this procedure until the entire asymmetric volume is sampled.

### Ligand restraints

Ligand restraints describing the covalent structures of non-standard OG, purine and AP site nucleotides were constructed with the *REEL* module of *Phenix*.[57] Attempts to use restraints from the ligand server or to build these ligand restraints by quantum mechanical calculations and energy minimization with eLBOW produced unnatural rings with, for example, unequal $C^{4'}$-$O^{4'}$ and $C^{1'}$-$O^{4'}$ bond lengths, or incorrectly attached hydrogen atoms. For this reason, restraints were edited manually starting with restraints for a standard adenine or guanine nucleotide and modifying atom names and adjusting parameters appropriate for atom substitutions. Values assigned for a subset of the torsion angles enforced specific sugar puckers, for example $C^{2'}$-*endo*. We updated restraints for the purine nucleotide (PRN), the OG nucleotide (8OG) and the abasic site (AAB, ORP) to match common sugar puckers. Additionally, we created alternate versions of the abasic site restraints to designate R or S chirality at the $C^{1'}$ position corresponding to *beta* and *alpha* anomers with the hydroxyl group pointing towards the 5' phosphate or the 3' phosphate. At first, we modeled the more common sugar puckers, $C^{2'}$-*endo* and $C^{3'}$*exo*, in combination with the *alpha* and *beta* stereoconfiguration to test if conformations could be excluded based on fit to electron density (**Figure S7A**). This approach excluded the $C^{3'}$-*endo beta* stereoisomer due to strong electron density map difference features (**Figure S7A**). Next, we expanded this analysis to include *alpha* and *beta* anomers for four different sugar puckers sampling across the pseudorotation landscape.[59] Of the different combinations, the *alpha* anomer fit electron density well, whereas the *beta* anomer yielded poorer matches, unreasonably high temperature factors for atom $O^{1'}$, lower correlation coefficients, and additional steric clashes (**Figure S7B**). Electron density calculated with phases from each of these eight possible structures consistently featured a bump on the 5-membered ring with equatorial disposition and extending towards the 5'-face indicating the model should be changed to the *alpha* anomer. This was the case even when phases were determined by the *beta* anomer structures, ruling out the possibility of phase bias. The simulated annealing composite omit map, which should be the least biased map, matched the shape for *alpha* anomers with an equatorial -OH group. Finally, we tested 10 different sugar puckers for the *alpha* anomer and selected $C^{4'}$-*endo* as the best conformation on the basis of fit to electron density and avoidance of steric clashes (**Figure S7C** and **Table S7**).

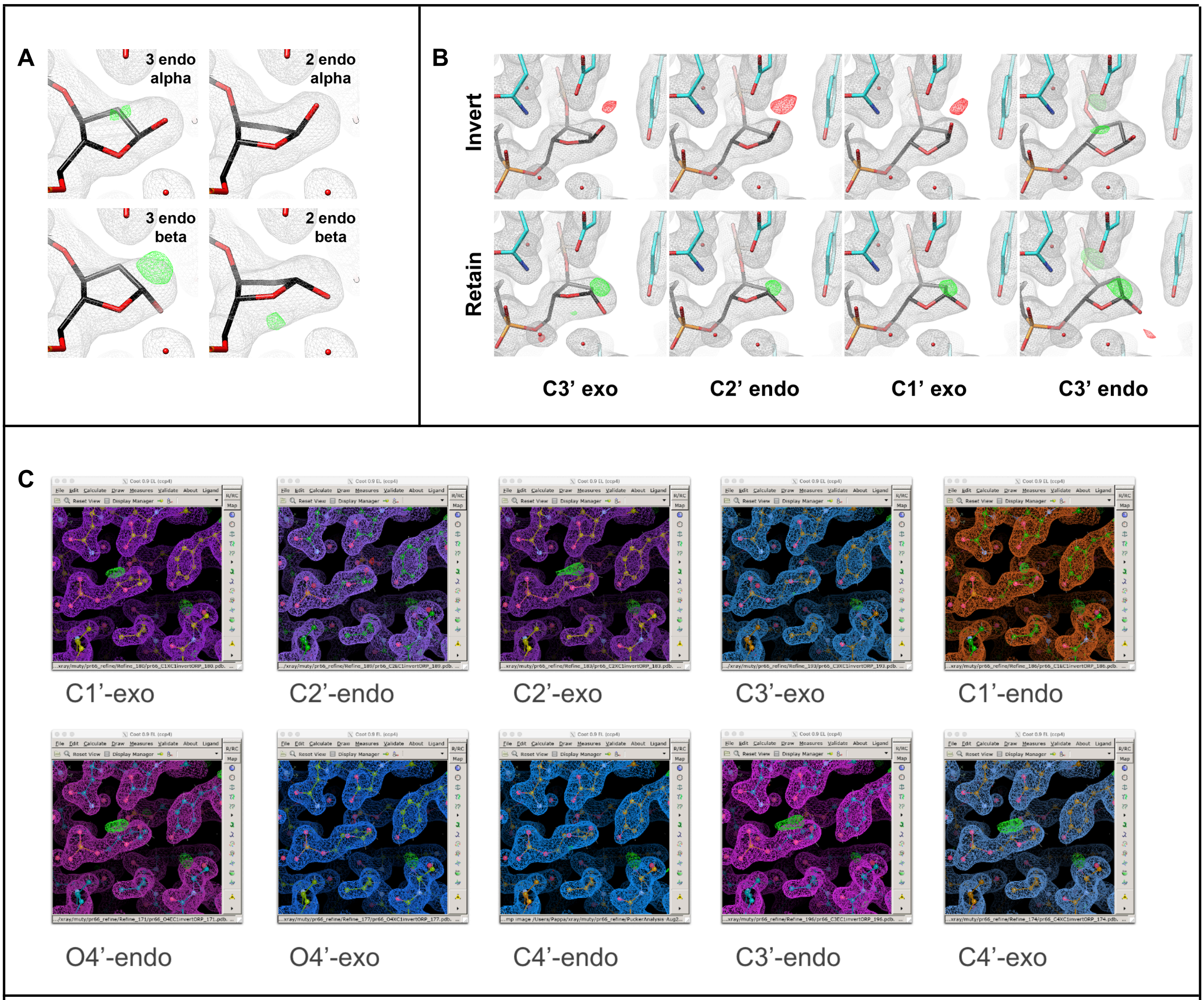

**Figure S7**. Comparison of different sugar puckers for *alpha* and *beta* anomers of the enzyme-generated AP site. Inversion of stereoconfiguration corresponds to the *alpha* anomer and retention generates the *beta* anomer. **A.** Initial tests with *alpha* and *beta* anomers and two common sugar puckers. **B.** More comprehensive tests for *alpha* and *beta* anomers and four sugar puckers. **C.** Testing 10 different sugar puckers for the *alpha* anomer. For torsion angles defining each sugar pucker see **Table S6** and **Figure S8**.

**Table S6**. Sugar pucker torsion angles (°)

| Angle | Atoms | C3'-endo | C4'-exo | O4'-endo | C1'-exo | C2'-endo | C3'-exo | C4'-endo | O4'-exo | C1'-endo | C2'-exo |
|---|---|---|---|---|---|---|---|---|---|---|---|
| 0 | C4',O4',C1',C2' | 2.8 | -18.0 | -26.7 | -35.8 | -19.3 | 1.6 | 18.0 | 26.7 | 35.8 | 19.3 |
| 1 | O4',C1',C2',C3' | -25.0 | 0.6 | 18.0 | 34.6 | 32.8 | 15.5 | -0.6 | -18.0 | -34.6 | -32.8 |
| 2 | C1',C2',C3',C4' | 35.9 | 16.5 | -0.6 | -20.2 | -33.1 | -24.3 | -16.5 | 0.6 | 20.2 | 33.1 |
| 3 | C2',C3',C4',O4' | -35.3 | -24.3 | -16.5 | -1.7 | 22.6 | 26.7 | 24.3 | 16.5 | 1.7 | -22.6 |
| 4 | C3',C4',O4',C1' | 20.5 | 26.7 | 24.3 | 23.1 | -2.3 | -18.0 | -26.7 | -24.3 | -23.1 | 2.3 |
| delta | C5',C4',C3',O3' | 84.8 | 92.4 | 102.5 | 124.8 | 145.2 | 164.2 | 143.7 | 132.5 | 118.9 | 95.8 |

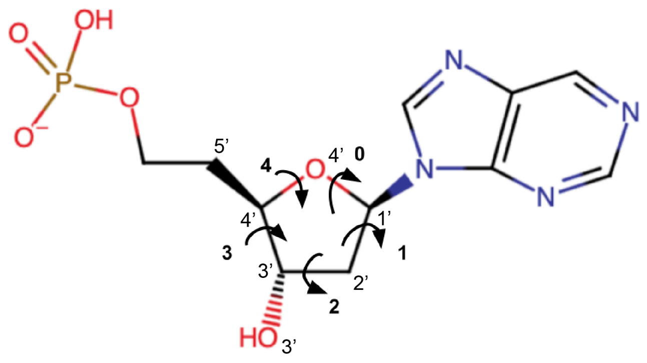

**Figure S8**. Torsion angles defining sugar pucker.

**Table S7**. Refinement outcomes for *alpha* anomer with different sugar puckers

| Metric | C3'-endo | C4'-exo | O4'-endo | C1'-exo | C2'-endo | C3'-exo | C4'-endo‡ | O4'-exo | C1'-endo | C2'-exo |
|---|---|---|---|---|---|---|---|---|---|---|
| Sum of steric overlaps | 0.963 | 0.940 | 0.514 | 0.912 | 0.421 | 0.878 | 0.403 | 0.495 | 0.514 | 0.974 |
| B value O1' (Å$^2$) | 38.10 | 37.06 | 36.79 | 38.02 | 39.86 | 38.93 | 38.77 | 38.28 | 39.45 | 39.33 |
| Map correlation* | 0.983 (0.91) | 0.983 (0.91) | 0.984 (0.92) | 0.985 (0.92) | 0.984 (0.92) | 0.984 (0.92) | 0.985 (0.92) | 0.986 (0.92) | 0.986 (0.92) | 0.985 (0.91) |
| \|Fo\| - \|Fc\| feature (e-/Å$^3$) | 0.3979 (+) | 0.4065 (+) | 0.3990 (+) | 0.3905 (+) | 0.3164 (+) | 0.2835 (+) | 0.2836 (+) | 0.2690 (+) | 0.2918 (+) | 0.3577 (+) |

* Correlation with |2Fo| - |Fc| (correlation with simulated annealing composite omit map)

‡ The final model was refined with C$^{4'}$-*endo* restraints (highlighted)
